# Evolutionary analysis supports variation in life history strategies between three foot-and-mouth-disease-virus serotypes

**DOI:** 10.64898/2026.08.18.745431

**Authors:** Avery Holmes, Eva Pérez-Martin, Simon Gubbins, Brianna Beechler, Anna Jolles, Roman Biek

**Affiliations:** School of Biodiversity, One Health & Veterinary Medicine, University of Glasgow, Glasgow, United Kingdom; The Pirbright Institute, Ash Road, Pirbright, Surrey GU24 0NF, UK; Department of Biomedical Sciences, Gary R Carlson MD College of Veterinary Medicine, Oregon State University, Corvallis, OR 97331, USA; Carlson College of Veterinary Medicine and Department of Integrative Biology, Oregon State University, Corvallis, OR, USA

## Abstract

Viruses have diverse life history strategies driven by variation in traits such as infectivity, transmission mode, and length and severity of infection that affect their epidemiology and evolution. While well documented among different species, life history and phenotypic variation among variants of the same virus species are less well understood. Foot-and-mouth-disease-virus (FMDV) is an ungulate-infecting picornavirus endemic to many regions, including Sub-Saharan Africa, where it circulates between wildlife and livestock in several serotypes. Recent work suggested that FMDV variants from the three Southern-African Territories serotypes exhibit different life history strategies, with these dynamics potentially causing distinct signatures in viral evolutionary rate, transmission among host species, and movement among regions. To investigate whether any effects of predicted effects occurred in natural settings, and whether these differences were shared with other strains within each serotype, this study used 716 published FMDV sequences (approximately 430bp) from 3 serotypes (SAT1, SAT2, and SAT3) to measure and compare evolutionary rates and transmission between regions and host types in Southern Africa. SAT1 had a slower rate of evolution consistent with a predicted more chronic infection strategy, and SAT2 had higher variability in evolutionary rates and some evidence of transmission from livestock to wildlife, suggesting livestock may play a part in persistence. SAT3 showed an expected intermediate phenotype but was challenging to validate due to small sample size. All SATs showed similar levels of transmission between regions. These results suggest that SAT1, SAT2, and SAT3 exhibit different transmission dynamics and evolutionary signatures, consistent with different life history strategies observed in their representative strains, such as more latency or a multi-host maintenance community.

## INTRODUCTION

Viruses exhibit considerable variation in their life histories (Holmes, 2009), analogous to the diversity of life history strategies seen in other organisms (Stearns, 1992). For example, viruses vary in host range and tissue specificity, as well as their timing and rate of replication, with some viruses causing short, acute infections, and others lingering for long periods in host cells and only replicating under certain conditions (Horner and Gale, 2013; Smith et al., 2019). Many herpes viruses lie dormant in the nervous system for years and only flare up when the host experiences physiological stress or old age (Gershon et al., 2015; Lu et al., 2021). These variations come with trade-offs, such as between virulence and transmissibility (Read and Harvey, 1993; Leggett et al., 2013), or between replication speed, genome size, and replication fidelity (Holmes, 2009). These trade-offs can cause ripple effects on viral ecology and infection dynamics and have significant implications for disease management (Bartlett, 1957; Baker et al., 2022). Ultimately, variation in life history traits is expected to be driven and reinforced by selective pressures varying across space and time, resulting in virus diversification and co-existence through niche partitioning (Mordecai et al., 2016). While life history and phenotypic variation among different viruses is well documented, how this plays out among variants of the same virus species is much less understood.

Although the effects of life history strategy differences between viruses may be challenging to observe directly, they may be inferred from evolutionary signatures in viral phylogenies. For example, latency is well understood to increase generation times and lower nucleotide substitution rates (Salemi et al., 1999; Vandamme et al., 2000). Therefore, latency is both a life history trait and a driver of measurable differences in viral evolution (Holmes, 2009). Other differences in life history characteristics, such as the R_0_ of the pathogen, may also cause differences in transmission that are detectible in phylogenetic analyses, such as through different rates of transmission between hosts or movement across the landscape. Certain evolutionary signatures of life history variation, such as the rate of molecular evolution, are well established when it comes to comparisons among virus species or families (Jenkins et al., 2002). In contrast, information at lower taxonomic scales, such as among variants or serotypes of the same virus species, is incomplete, limiting our understanding of how genetic and phenotypic diversity in viruses is generated and maintained.

Foot and mouth disease virus (FMDV) offers an intriguing system to examine the diversity of life history strategies and evolutionary dynamics in closely related variants of the same virus. FMDV is a single-stranded RNA virus in the family *Picornaviridae* that infects even-toed ungulate animals such as sheep, pigs, and cattle (Jamal and Belsham, 2013). It causes foot-and-mouth disease (FMD), an economically significant livestock disease that remains endemic in many parts of world and has caused recent outbreaks in non-endemic areas, including Germany and Hungary (European Comission, 2025a, 2025b). Endemic regions include sub-Saharan Africa, where FMDV circulates in five main subtypes: South African Territories (SAT) types 1, 2, and 3, which infect African buffalo (*Syncerus caffer*) and occasionally cattle, and serotypes A and O, which are predominantly maintained by cattle (Casey-Bryars et al., 2018). Although FMDV infection rarely results in host death, it causes high morbidity, leading to lowered meat and milk production, abortion of calves, decreased draught power, and lost cash revenue from sales that can be devastating to subsistence farming communities (Knight-Jones and Rushton, 2013). Control efforts are hampered by FMDV being one of the most infectious pathogens known to science (Jolles et al., 2021).

FMDV serotypes SAT1, 2, and 3 circulate simultaneously in African buffalo, suggesting that all three serotypes have evolved life history strategies compatible with buffalo population ecology (Hampson and Haydon, 2021), which may include the ability to cause chronic carrier infections, exploiting waning host immunity, or fast antigenic evolution (Jolles et al., 2021). In fact, various strains of FMDV, including type O and all three SATs, have been recovered from animals long after clinical signs have passed, suggesting that chronic infections occur (Bertram et al., 2018; Cortey et al., 2019; Jolles et al., 2021). Additionally, three FMDV strains each representing one of the three South African FMDV types exhibit different levels of persistence within hosts (Jolles et al., 2021), and SAT2 has been detected outside of Sub-Saharan Africa on multiple occasions whereas other SATs have not (Hall et al., 2013; WOAH and FAO, 2023), potentially suggesting variable transmission dynamics between SATs. This suggests that some FMDV strains have evolved different strategies to persist in buffalo populations. However, it remains untested if and how life history differences among FMDV strains have shaped their molecular evolution and epidemiology, and whether these differences are present within other strains of the same serotype.

A long history of FMDV testing and research in southern Africa makes this area, specifically the countries of South Africa, Botswana, Zimbabwe, and Mozambique, an ideal study system for these questions. Sampling has been especially focused in and around Kruger National Park (KNP), which is located near the South African North-East border with Mozambique. KNP has been a hot-spot for FMDV research, since the park and its African buffalo population are thought to be a reservoir and source of FMDV within the larger geographic area, with livestock and neighbouring regions likely acting as sinks (Jori and Etter, 2016). The wealth of genetic information collected over multiple decades from KNP and Southern Africa offers a unique opportunity to quantify the evolution, inter-regional transmission, and between-host transmission dynamics of SAT 1, 2 and 3 and to test for viral evolutionary signals consistent with life history strategy variation. Specifically, we sought to determine whether the three serotypes show differences with respect to their:

1. evolutionary rate, in terms of the mean rate and its variability - we predicted a lower evolutionary rate for SAT1 compared to SAT2 and SAT3 due to higher propensity for within-host persistence (Jolles et al. 2021);
2. patterns of viral movement among regions and contribution of KNP as a source, with SAT2 expected to show evidence of more long-range movement (Hall et al., 2013; WOAH and FAO, 2023);
3. levels of transmission between wildlife and livestock and ability to persist in livestock populations, with SAT2 expected to show more transmission within livestock (Dyason, 2010; Fana et al., 2021; Hall et al., 2013).

## METHODS

### Collection of sequence data

All available FMDV sequences for the VP1 gene, in full genomes, and their metadata were retrieved from GenBank using the Rentrez package in R (version 4.2.2) (Winter, 2017; R Core Team, 2023) in October 2022 using the search terms “FMDV”, “South Africa”, “Botswana”, “Zimbabwe”, and “Mozambique”. The search terms “Eswatini” and “Lesotho” were also included, but no sequences were available from these countries. Some sequences collected from other regions were included in the results of the search due to having been published by an institution in one of the target countries; these sequences were removed. Metadata collected included serotype, region, host, and date. If any metadata was missing, the original publication from the sequence was consulted and the data updated manually. Sequences identified as a different serotype other than SAT1, 2, and 3 (such as A or O) were removed. Six sequences from SAT1 without sampling date information were excluded. Most of the available sequences originated from wildlife due to ongoing active surveillance efforts, especially in KNP. In contrast, livestock were generally sampled in the context of an outbreak.

All sequences were aligned using MUSCLE alignment (Edgar, 2004) in Geneious (version 2022.1.1) (http://www.geneious.com), trimmed to include only nucleotides for VP1, and a maximum likelihood tree of all sequences was created using MEGA11 (version 11.0.8) (Tamura et al., 2021) using a GTR+G+I evolutionary model. Sequences with missing serotype information were then assigned a SAT based on which clade they were placed in.

Following serotype assignment, a new separate alignment was created for each SAT and trimmed to VP1, including any adjacent nucleotides that were included in >75% of the available sequences. These alignments (SAT1: n= 269, bp= 436; SAT2: n= 315, bp= 423; SAT3: n= 132, bp= 465) were used for all further analyses.

### Organisation of metadata

The time of sequence collection was coded with the day, month, and year. When only the month of collection was available, the 15^th^ of each month was assigned, and if only the year was available the first of July of that year was assigned. Location was coded as a discrete trait with five states: Botswana (BOT), Kruger National Park (KNP) South African Republic (SAR), Mozambique (MOZ), and Zimbabwe (ZIM) (Table 1). Treating KNP as a separate location, was immediately apparent from isolate names, allowed for better spatial resolution and was justified by a large proportion of available sequences originating from KNP. Although KNP borders other parks in neighbouring countries as part of the Great Limpopo Transfrontier Conservation Area (Figure 1), metadata was not sufficient to identify whether the samples from these neighbouring countries were from within this conservation area or not.

**Figure 1:**
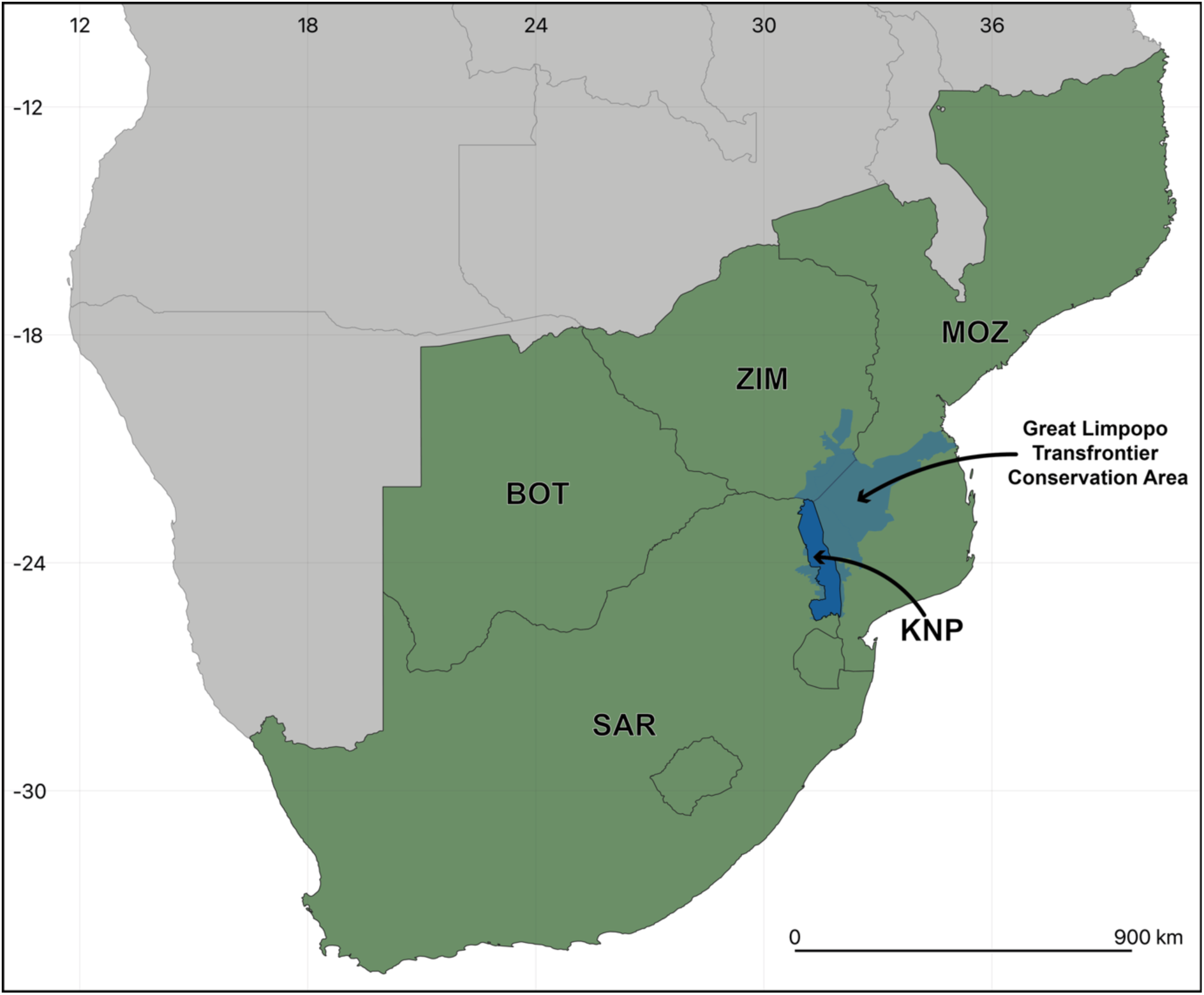
**Geographic regions included in this study**, with the Kruger National Park (KNP) highlighted in dark blue and the Great Limpopo Transfrontier Conservation Area highlighted in paler blue. MOZ = Mozambique, ZIM = Zimbabwe, BOT = Botswana, SAR = South African Republic.

**Table 1:** Number of sequences for each FMDV serotype in each geographic region. BOT = Botswana, KNP = Kruger National Park, SAR = South African Republic, MOZ = Mozambique, ZIM = Zimbabwe.

| Serotype | BOT | KNP | SAR | MOZ | ZIM | Total |
| --- | --- | --- | --- | --- | --- | --- |
| SAT1 | 23 | 63 | 39 | 6 | 138 | 269 |
| SAT2 | 18 | 93 | 71 | 13 | 120 | 315 |
| SAT3 | 10 | 30 | 6 | 3 | 83 | 132 |

Although most hosts were either buffalo or cattle, some viruses were isolated from other wildlife species, including impalas and kudus, and domestic species, including sheep. These poorly represented species were considered unlikely to play a significant role in FMDV transmission. Therefore, their sequences were included, and ‘host type’ was coded as ‘wildlife’ (SAT1 = 211, SAT2 = 171, SAT3 = 103) or ‘livestock’ (SAT1 = 47, SAT2 = 132, SAT3 = 24) as another discrete trait. Host information was unavailable for 35 sequences, and these were coded as unknown (SAT1 = 17, SAT2 = 12, SAT3 = 6), with these states subsequently inferred in the model.

### Assessment of a clock-like signal and analysis with BEAST

To investigate whether applying molecular clock models to the sequences would be appropriate, a maximum likelihood tree was made with MEGA11 for each SAT and analysed for clock-like signal with TempEst (version 1.5.3) (Rambaut et al., 2016). To gain a clearer picture of any variability in the clock signal for each SAT, the two or three largest clades in each tree were also analysed independently with TempEst. This analysis showed a clear clock-like signal for SAT1 and SAT2, but a much weaker and less consistent signal for SAT3 (Table S1, supplementary figures S1-S12). To verify these results, a Bayesian evaluation of temporal signal (BETS) was performed (Duchene et al., 2020). For each subtree of each SAT, two models were run in BEAST (version 1.10.4)(Suchard et al., 2018), one with all tip dates set to 0 and a fixed clock rate (isochronous model), and one with tip dates included and a relaxed clock rate (heterochronous model). For all models, a constant population size was assumed, a SRD06 evolutionary model was used (Shapiro et al., 2006), and the log marginal likelihood was estimated with generalised stepping stone sampling using the product of exponential distributions and an underlying beta distribution (Duchene et al., 2020). Each model was run for 100k states. A Bayes factor test was used to compare the likelihoods of each model for each subtree, and the heterochronous model was preferred in all cases, with Bayes factor values ranging from 4.7 to 247, indicating very strong support for a clock signal in all SATs (Table S2.)

Evolutionary rate, rates of transmission between regions, and rates of transmission between host types were estimated in BEAST (version 1.10.4) (Suchard et al., 2018) using a SRD06 evolutionary model (Shapiro et al., 2006), a relaxed clock rate with a lognormal distribution, and a Bayesian SkyGrid demographic model to allow flexibility in population history and account for variation in sampling effort (Hill and Baele, 2019). The rates of evolution, regional transmission, and host type association were estimated together in one model by discrete trait analysis (DTA). This model estimated both the instantaneous rate of regional and host type change and also the number of Markov jumps for region and host type, which indicates the number of discrete state transmission events observed in the data (Minin and Suchard, 2007, 2008; Drummond et al., 2012). Markov jump estimates offer a more objective description of the rates of change compared to instantaneous rate estimates, which are based on the assumption that the dynamics of state change are consistent across the entire evolutionary history represented by the tree. Three analyses consisting of 200 million states each were run independently for each SAT and were compared to confirm convergence. The burn-in was lengthened as necessary in cases where visual inspection suggested that parameter estimate convergence had not been reached by the standard cut-off of 10% of the states. The estimates from all three runs were combined to provide the final results. Because the number of sequences for each SAT varied, Markov jump counts were presented both raw and corrected via dividing by the number of sequences to account for the fact that more jumps can be observed in larger datasets. Maximum clade credibility trees were made for each SAT with TreeAnnotator v.1.10.4 (Suchard et al., 2018) and visualised with ggtree (Yu, 2020).

### Post-hoc tests of host type associations with evolutionary rates and regional transmission events

Due to a high level of variability in the evolutionary rate of SAT2 across different branches, the distributions of SAT2 evolutionary rates in livestock and wildlife from the DTA MCC tree were compared to examine whether the virus evolves at different rates in different host types. Additionally, to test whether regional movement was facilitated more by either livestock or wildlife, the number of transitions from wildlife to wildlife, wildlife to livestock, livestock to wildlife, and livestock to livestock in the trees were counted and a Fisher’s exact test was run to assess whether inter-regional transitions were over-represented in that host-transmission type in relation to the frequency of that host-transition in the tree. Within each SAT, p-values were corrected for multiple testing (Benjamini and Hochberg, 1995).

### Validation of discrete trait analysis host transmission results by structured coalescent approximation

Because DTA is known to be vulnerable to bias when sampling between groups has been uneven (Layan et al., 2023), a structured coalescent approximation approach in BEAST v.2.7.6 (Bouckaert et al., 2019) using MASCOT v.3.0.7 (Müller et al., 2024) was used to validate the transmission rate estimates between wildlife and livestock for each SAT, as this method makes different assumptions and is more robust to sampling imbalance. Samples with missing host information were removed, since MASCOT does not infer their host association. Since the SRD06 model used for DTA was not available for MASCOT, the evolutionary model was set to TIM2+R4 after model selection with IQ-Tree (Minh et al., 2020), and the evolutionary clock was set to optimised relaxed clock. To allow for changing host demographics, host population priors were set to Skyline for both wildlife and livestock populations, with all N_e_ priors set to a normal distribution with µ = 0 and σ = 1, as recommended uninformative starting priors. Each model was run three times for 200 million states each. Some of the SAT1 runs produced negative infinity and a sharp collapse in likelihood, so additional runs were set up for SAT1 until three successful runs had been obtained. Only runs that ran successfully to 200 million states were included in the results. The first 10% of states were removed from all runs as burn-in. Maximum clade credibility trees for each SAT were made with Tree Annotator v.2.7.6 (Bouckaert et al., 2019) and visualised with ggtree (Yu, 2020). As with the DTA results, Markov jump counts are presented both raw and corrected via dividing by the number of sequences to account for differing sample sizes between SATs.

## RESULTS

### Evolutionary rates

The BEAST estimates for evolutionary rate showed similar mean values for all SATs, with all 95% highest posterior density intervals (HPDI) estimates overlapping (SAT1 HPDI = 2.90×10^−3^ - 4.71×10^−3^ substitutions/site/year (s/s/y); SAT2 HPDI = 3.23×10^−3^ - 5.33×10^−3^ s/s/y; SAT3 HPDI = 3.04×10^−3^ - 5.65×10^−3^ s/s/y), although the estimates for SAT3 had a low effective sample size and struggled to definitively converge on a range of values (Figure 2 A). SAT2 showed a considerably higher coefficient of variation, meaning that the evolutionary rate varied much more along different branches of the tree compared to SAT1 or 3 (SAT1: HPDI = 4.50×10^−1^ - 7.70×10^−1^. SAT2: HPDI = 8.05×10^−1^ - 1.49. SAT3: HPDI = 4.53×10^−1^ - 9.60×10^−1^) (Figure 2 B).

**Figure 2:**
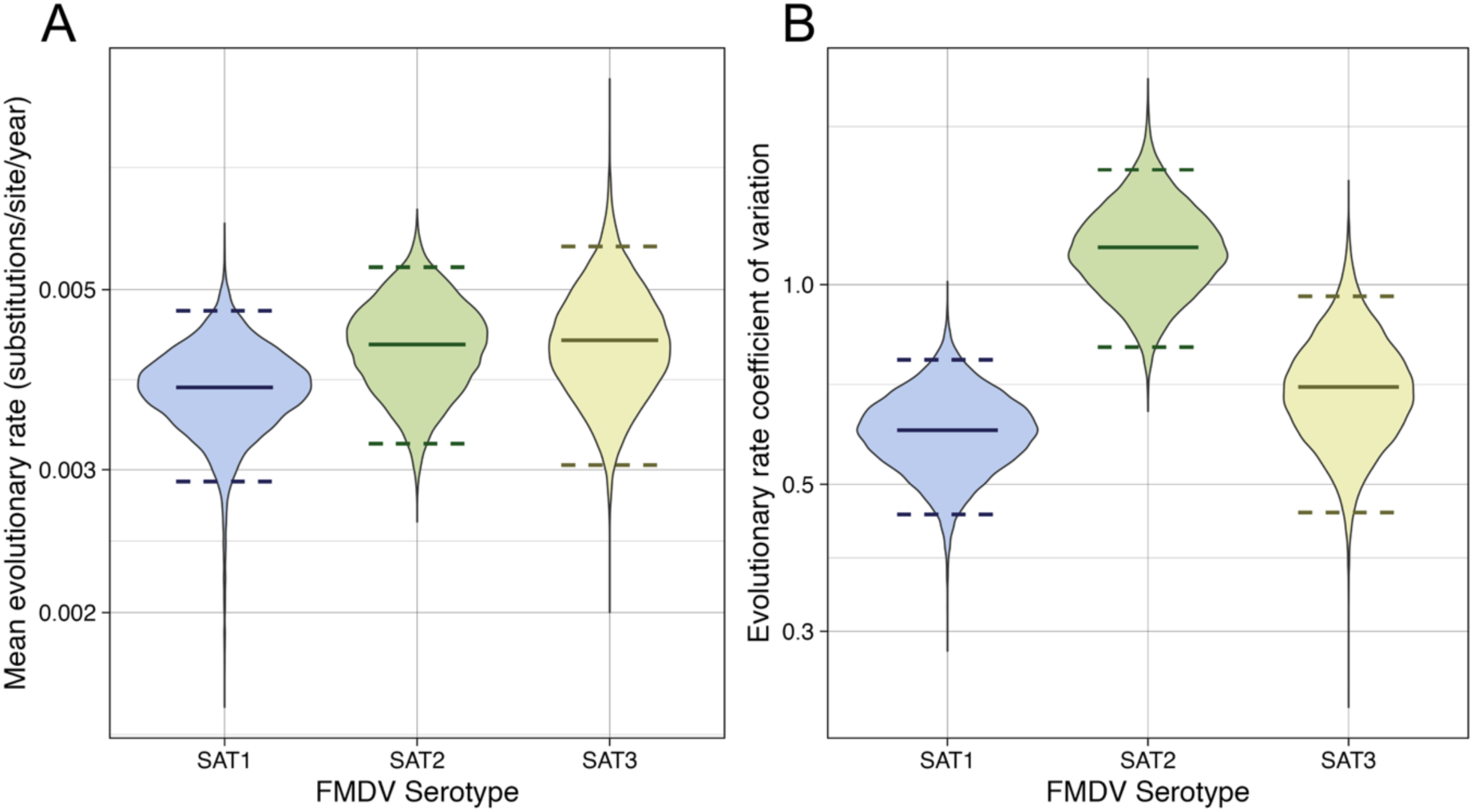
Evolutionary rate estimates for FMDV serotypes SAT1, 2 and 3. Estimates obtained in BEASTv1.10.4. Solid lines indicate mean values, dashed lines indicate 95% highest posterior density interval (HPDI). A) Mean substitution rate per site per year, with SAT1 evolving somewhat slower than SAT2 or SAT3, although the rate for SAT3 did not converge well and is therefore slightly less reliable. B) Evolutionary rate coefficient of variation, with SAT2 considerably more variable than SAT1, and SAT3 having intermediate variability with ranges more similar to SAT1 than SAT2.

We asked whether the variation in evolutionary rates seen within the trees (an especially in SAT2) could be a result of different evolutionary rates in different host types. The distributions of evolutionary branch rates of branches inferred to be associated with livestock and wildlife broadly overlapped in all SATs, suggesting that FMDV did not consistently evolve more quickly in one type of host (Figure 3). However, the range in branch rates for livestock was smaller than for wildlife for SAT1 and SAT2, suggesting that FMDV of these serotypes may evolve more consistently in livestock. A small proportion of branches associated with wildlife evolved very quickly, at rates of more than 1×10^−2^ substitutions per site per year.

**Figure 3:**
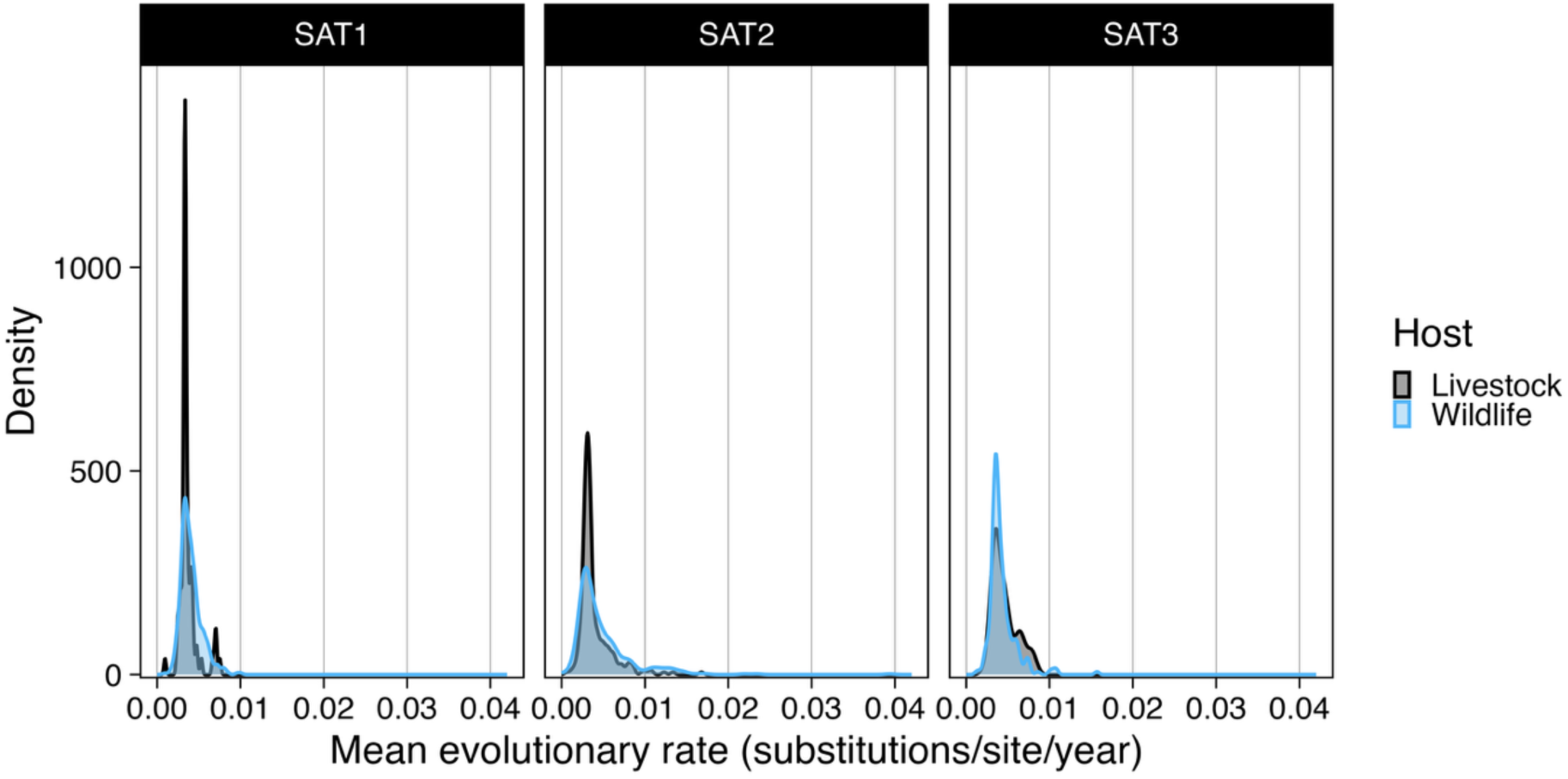
SAT2 Evolutionary rate distributions for FMDV branches from livestock and wildlife. All SATs evolved at generally similar rates in both livestock and wildlife. However, the wildlife-associated branches show a wide shoulder in SATs 1 and 2, with some branches evolving at a rate of over 1.0×10^−2^, especially in SAT2

### Viral movement between regions

All SATs had overlapping rates of inter-regional movement (SAT1 HPDI = 2.40×10^−3^ - 7.47×10^−3^ events/year (e/y); SAT2 HPDI = 3.27×10^−3^ - 9.11×10^−3^ e/y; SAT3 HPDI = 2.65×10^−3^ - 1.06×10^−2^ e/y) (Figure 4A). Additionally, the number of observed transition events were similar between SATs after correction for sample size (Table 2). The coefficient of variation for viral movement was very similar for all serotypes (SAT1 HPDI = 2.98×10^−5^ - 2.92; SAT2 HPDI = 1.49×10^−4^ - 2.02. SAT3: HPDI = 2.74×10^−5^ - 2.45) (Figure 4B).

**Figure 4:**
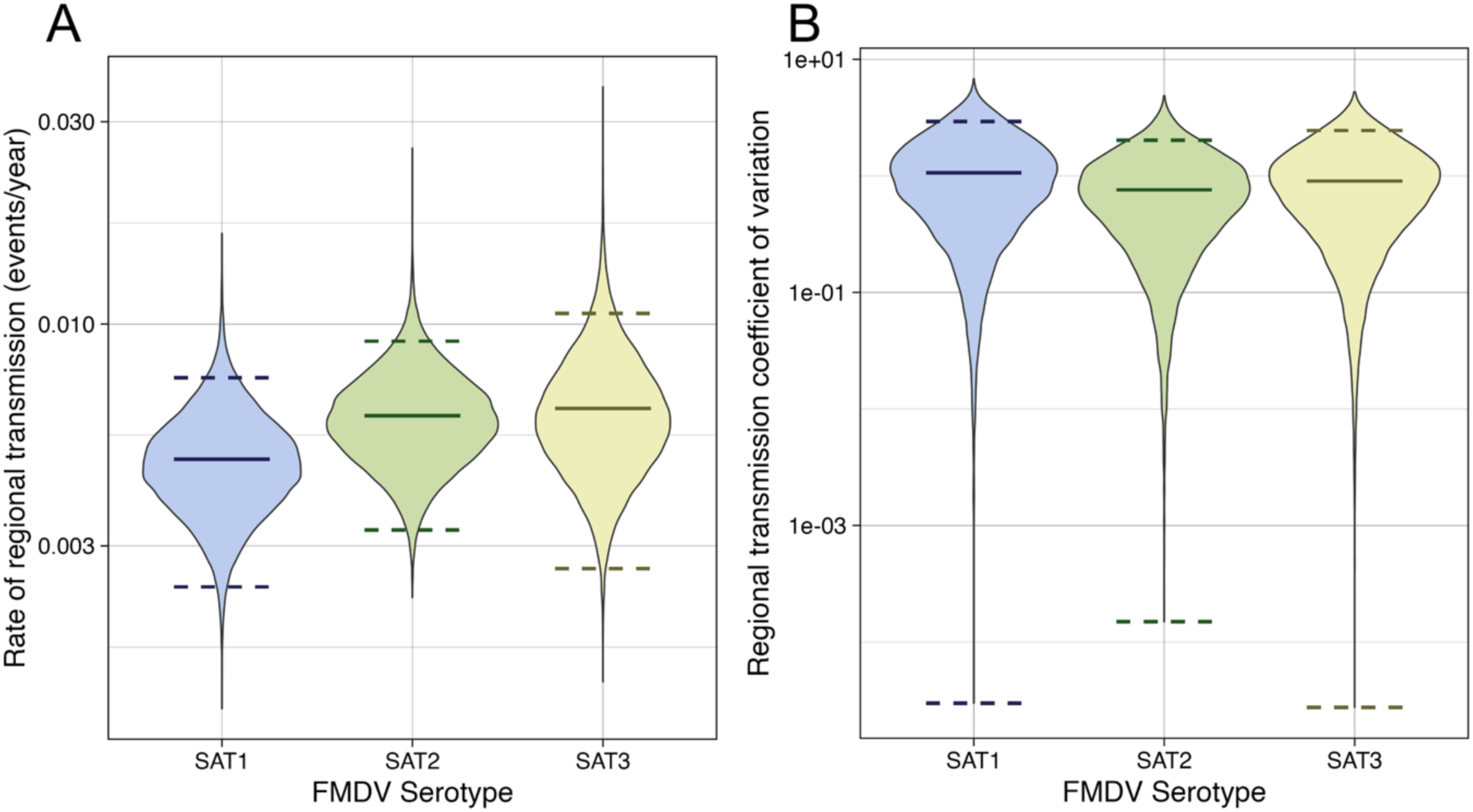
Estimated rates of movement between regions for SAT1, 2, and 3. Estimates obtained in BEASTv1.10.4. Solid lines indicate mean values, and dashed lines indicate 95% highest posterior density interval (HPDI). A) Rate of inter-region transmission for all SATs, with SAT1 showing slightly lower estimates than SAT2, and the estimates for SAT3 being more variable with a wider HPDI. B) Coefficient of variation in inter-region transmission for all SATs, with similar estimates for all SATs, but a slightly higher mean for SAT1. All HPDIs include the minimum estimate.

**Table 2:** Discrete between-region movement estimates inferred from each tree, corrected by number of sequences for each SAT to account for larger sample sizes leading to more opportunities to observe movement events, facilitating direct comparison between SATs.

| SAT | Region transitions | HPDI | N sequences | Transitions/Sequence |
| --- | --- | --- | --- | --- |
| SAT1 | 26.614 | 22-31 | 269 | 0.099 |
| SAT2 | 31.155 | 27-36 | 315 | 0.099 |
| SAT3 | 19.228 | 15-24 | 132 | 0.146 |

The rates of inter-region transmission were similar between SATs, and for all three the greatest transmission link was from KNP to some other region, often South Africa or Zimbabwe (Figure 5). However, although KNP was more often a source of virus than a sink, transmission back into the park was not uncommon, with notable transmission into the park from South Africa in SAT1 and Zimbabwe in SAT3. Additionally, viral exchange between other regions not mediated by KNP, such as between Zimbabwe and Botswana in SAT2, also showed evidence of considerable transmission.

**Figure 5:**
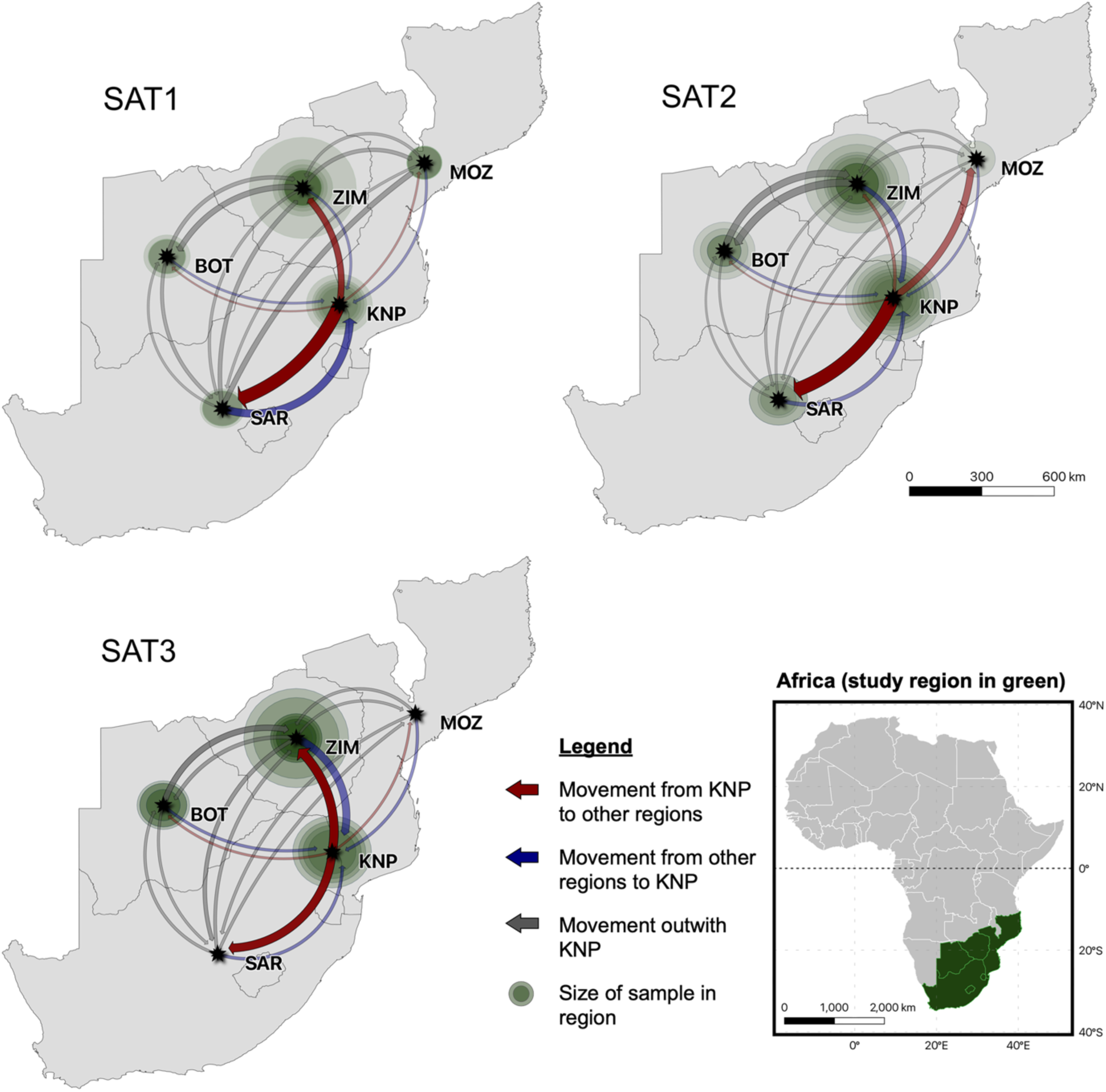
Transmission of FMDV between regions. Kruger National Park (KNP), Mozambique (MOZ), Zimbabwe (ZIM), Botswana (BOT), and South Africa (SAR). Red arrows indicate viral transmission out of KNP. Blue arrows indicate viral transmission into KNP. Grey arrows indicate transmission between other regions. Each arrow’s thickness indicates total transmission rate, and each arrow’s opacity represents transmission rate relative to other arrows on the same map. Black points represent the centre of each region and may not be accurate to the sampling location, which is often unavailable. The green circle around each region point indicates the number of sequences included for that serotype in that region. Inset: Africa with the countries included in this study highlighted in green.

We asked whether inter-region transmission events were associated with types of between-host transmission events (e.g., wildlife to livestock and wildlife to wildlife transmission events) compared to the expected frequencies of based on the numbers of each host transmission type. In SAT1, wildlife-to-wildlife transmission events showed a higher rate of inter-region transmission events than expected, whereas wildlife-to-livestock showed a lower rate than expected (Table 3). A lower rate from wildlife to livestock was also observed in SAT2, which, although only marginally significant, was consistent with the pattern in SAT1. All other SATs showed expected levels of inter-region transmission in each host transmission type.

**Table 3:** Tests of association between host transition type and inter-region transition events. Fisher’s exact tests with odds ratios and p-values corrected for multiple testing are presented for each SAT and each host transition type. W -> W = Wildlife to wildlife transition, W -> L = Wildlife to livestock transition, L -> L = Livestock to livestock transition, L -> W = Livestock to wildlife transition.

| SAT | N | N Livestock | N Wildlife | Shift type | Count | P | Odds ratio | 95% CI |
| --- | --- | --- | --- | --- | --- | --- | --- | --- |
| SAT1 | 537 | 74 | 463 | W -> W | 12 | **0.02** | 5.86 | 1.22 - 39.38 |
|  |  |  |  | W -> L | 10 | **<0.01** | 0.06 | 0.00 - 0.53 |
|  |  |  |  | L -> L | 1 | 1.00 | 2.06 | 0.10 - 129.03 |
|  |  |  |  | L -> W | 0 | 1.00 | 0.00 | 0.00 - Inf |
| SAT2 | 629 | 255 | 374 | W -> W | 16 | 1.00 | 1.09 | 0.33 - 3.64 |
|  |  |  |  | W -> L | 7 | **0.05** | 0.12 | 0.00 - 1.05 |
|  |  |  |  | L -> L | 5 | 0.14 | 2.66 | 0.68 - 11.86 |
|  |  |  |  | L -> W | 0 | 1.00 | 0.00 | 0.00 - Inf |
| SAT3 | 263 | 42 | 221 | W -> W | 14 | 1.00 | 0.79 | 0.06 - 6.40 |
|  |  |  |  | W -> L | 1 | 1.00 | 0.61 | 0.01 - 50.21 |
|  |  |  |  | L -> L | 1 | 1.00 | 1.93 | 0.14 - 109.39 |
|  |  |  |  | L -> W | 0 | 1.00 | 0.00 | 0.00 - Inf |

### Transmission between host types

MCC trees generated from both DTA and MASCOT analyses for each SAT showed a similar topology, and visual examination of both sets of trees illustrated sequences isolated from livestock generally located on the tips and recent internal nodes, whereas the deeper nodes (from times when disease control around KNP was more rigorous and South Africa was officially FMDV-free) were entirely assigned to wildlife. SAT1 MASCOT results had an additional alternative convergence state in which the root was assigned to livestock and wildlife were indicated as spillover hosts. However, this state required that the viral effective population size in livestock be considerably larger than in wildlife, whereas for all other runs in other SATs, the effective population size in wildlife was inferred to be much larger. Since the latter result was also much more consistent with current knowledge of FMDV epidemiology, only results for which wildlife had the larger effective population size estimate are presented here.

The trees for SAT1 showed wildlife branches were considerably more common than livestock branches, indicative of the virus being maintained by wildlife hosts. Association with livestock tended to be seen on the tips and was underrepresented deeper in the tree. Host type shifts appeared to be predominantly from wildlife to livestock, although DTA and structured coalescent did not always agree on the transmission events happening in the same place – for example, at the parent or daughter node (Figure 6). A few outbreaks in livestock could be seen in closely related taxa with very short branch lengths.

**Figure 6.**
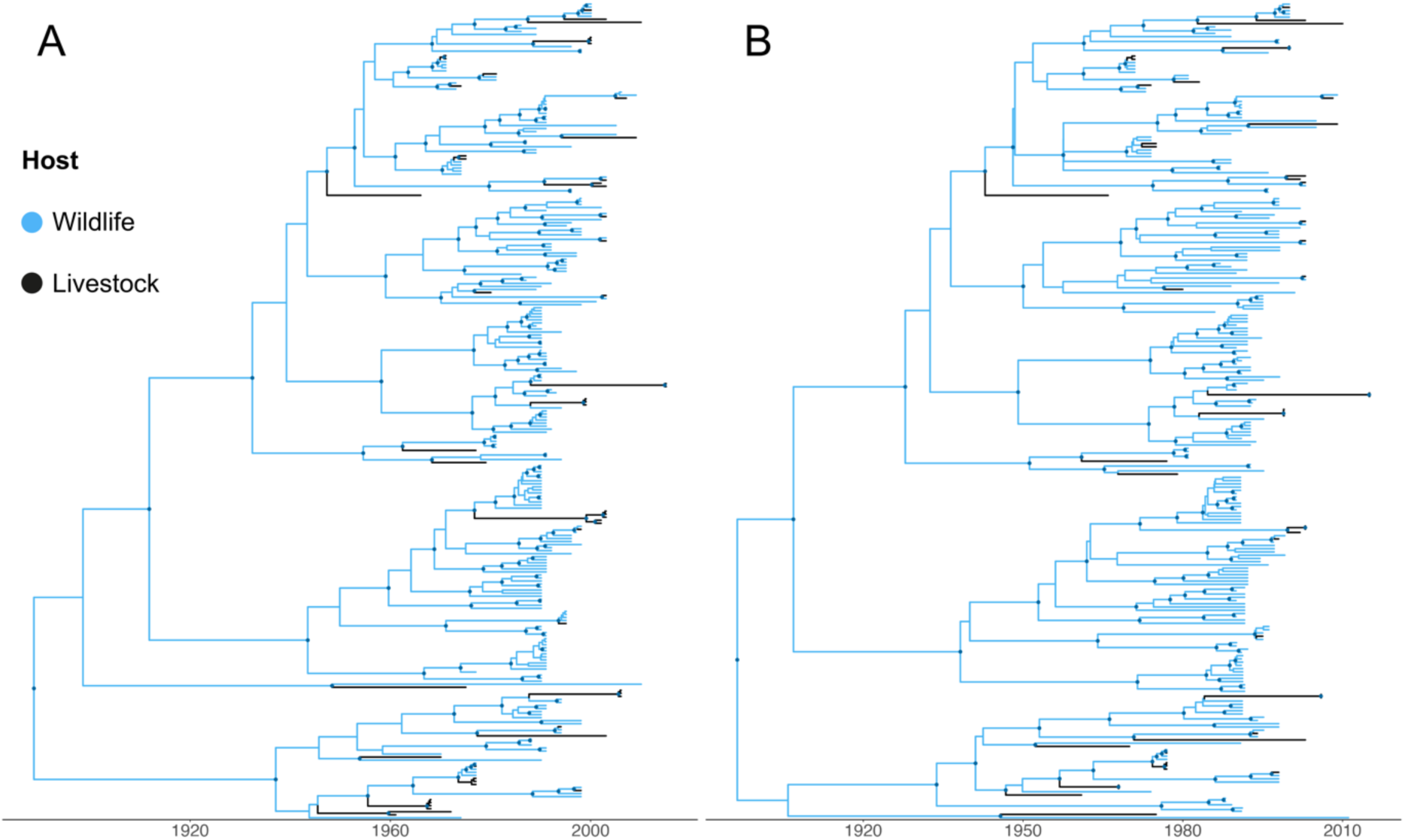
**SAT1 maximum clade credibility trees,** with nodes with posterior support of 0.8 or more indicated by darker dots. Both trees show livestock branches near the tips of the tree, with all deep branches assigned to wildlife. Some small outbreaks of transmission within livestock are visible. A) Tree generated from discrete trait analysis in BEAST. B) Tree generated from structured coalescent approximation in MASCOT using BEAST2.

Similarly to SAT1, the SAT2 MCC trees showed livestock were represented mostly at the tips of the tree, with most internal branches assigned to wildlife (Figure 7). Some livestock tips clumped together on short branches, which is consistent with single introductions that led to outbreaks of sustained transmission in livestock, which were larger and more common (mean livestock tips per wildlife-livestock transmission event: 5.3 DTA, 3.6 MASCOT, median: 2.5 DTA, 2 MASCOT, maximum: 26 both DTA and MASCOT) than those observed in SAT1 (mean: 1.6 DTA, 1.4 MASCOT, median: 1 both DTA and MASCOT, maximum: 5 DTA, 3 MASCOT) or SAT3 (mean: 2.3 DTA, 1.8 MASCOT, median: 1.5 DTA, 1 MASCOT, maximum: 7 DTA, 4 MASCOT). Also, unlike other trees, the SAT2 discrete trait analysis tree clearly showed seven instances of transmission from livestock to wildlife. However, these transmission events were not supported in the MASCOT analysis – for the equivalent sections of the MCC tree, the SAT2 viruses were inferred to have remained associated with wildlife, resulting in only transmission from wildlife to livestock being supported (Fig 6 B).

**Figure 7:**
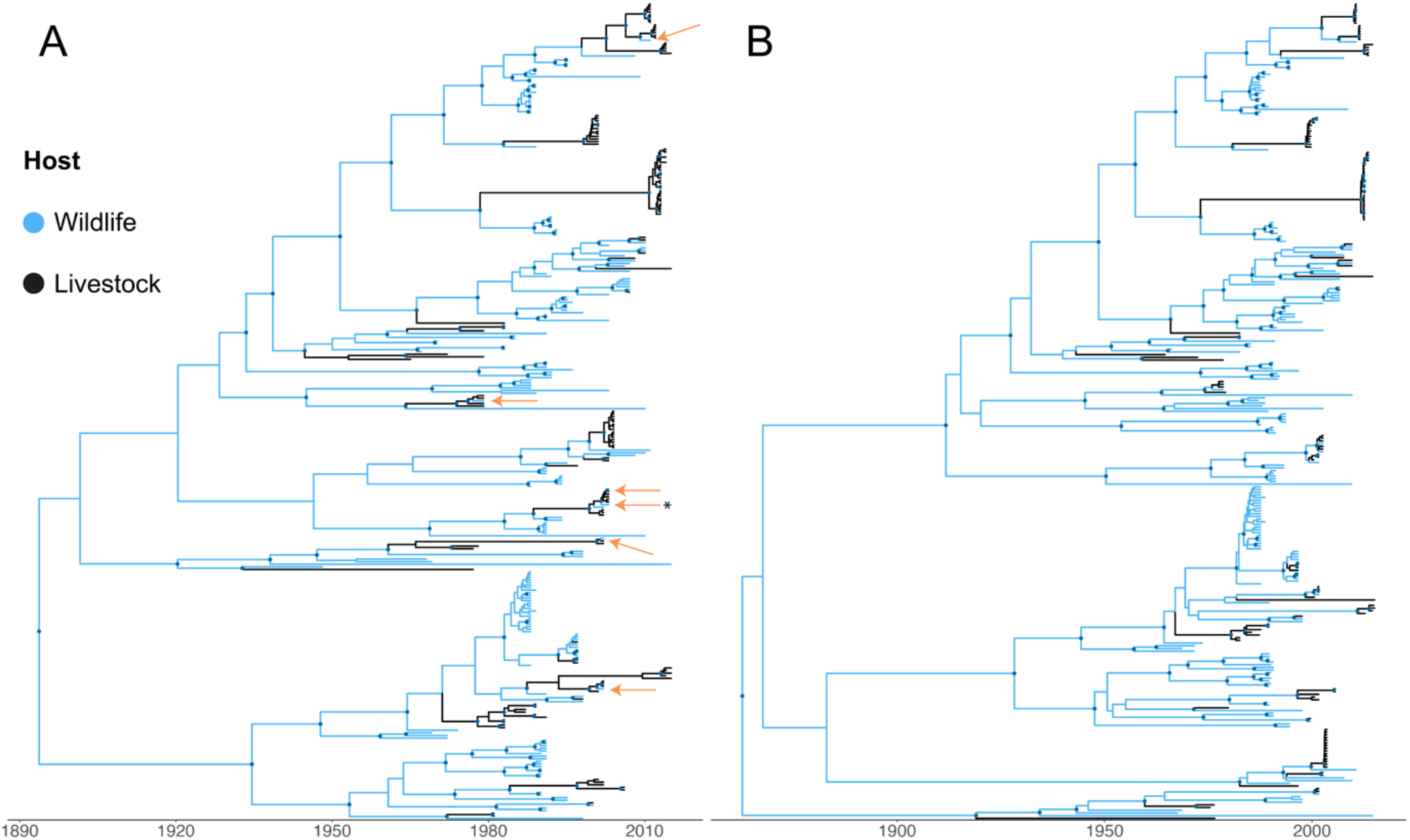
**SAT2 maximum clade credibility trees,** with nodes with posterior support of 0.8 or more indicated by darker dots. Both trees show livestock branches occur near the tips of the tree, with all deep branches assigned to wildlife. Several outbreaks involving transmission within livestock are visible in both trees, with some of these of a considerable size. A) Tree generated from discrete trait analysis in BEAST, with inferred livestock to wildlife transmission events indicated with orange arrows, one of which points to two transmission events close within the tree (marked with an asterisk). B) Tree generated from structured coalescent approximation in MASCOT using BEAST2, with the transmission events from livestock to wildlife in the BEAST DTA tree not supported.

The tree for SAT3 also showed livestock represented only at the ends of branches, and livestock were underrepresented deep in the tree (Figure 8). Similarly to SAT1 but unlike SAT2, the SAT3 tree showed limited evidence for sustained transmission in livestock.

**Figure 8:**
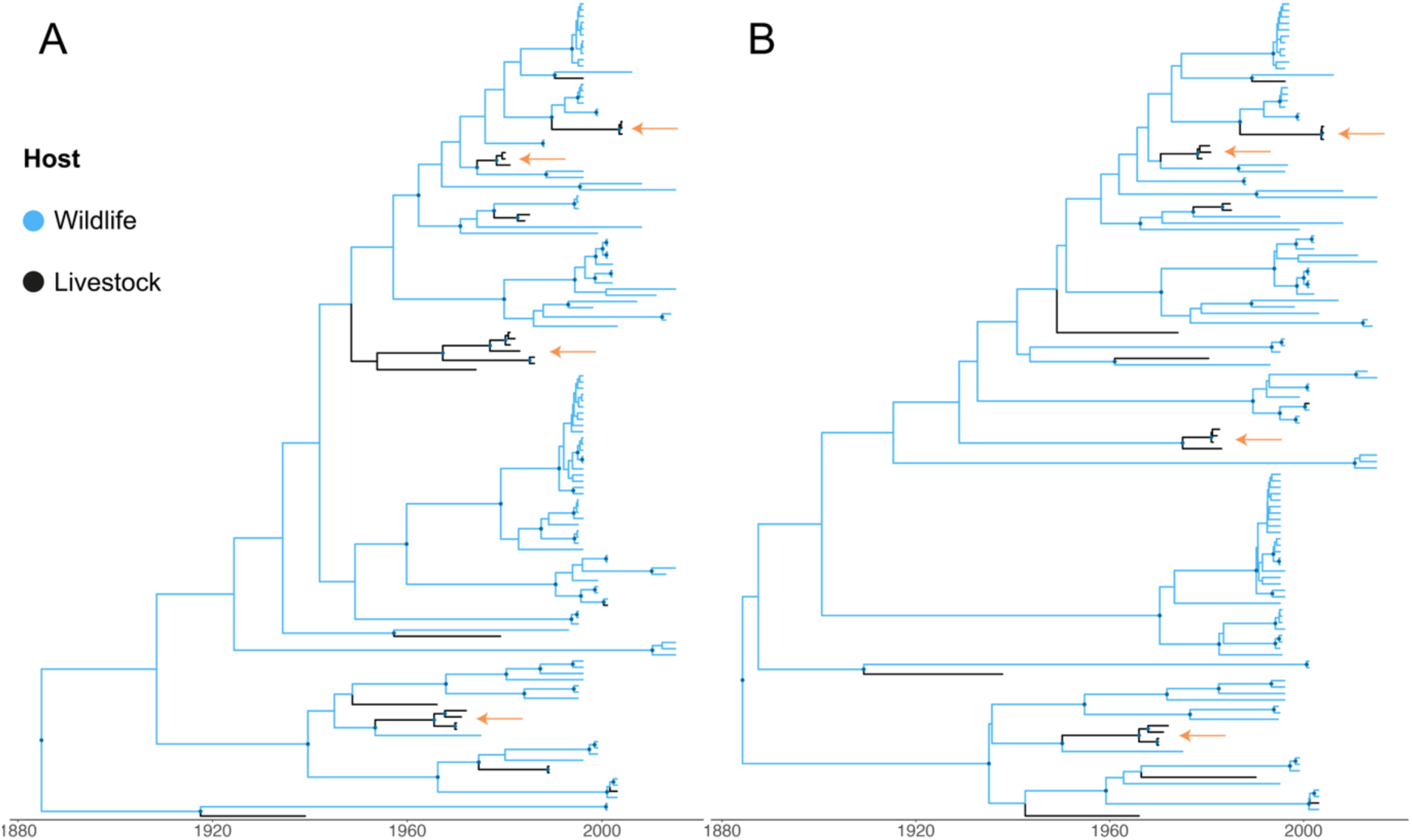
**SAT3 maximum clade credibility trees,** with nodes with posterior support of 0.8 or more indicated by darker dots. Both trees show livestock branches occur near the tips of the tree, with all deep branches assigned to wildlife. Some small outbreaks of transmission within livestock are visible in both trees (marked with orange arrows). A) Tree generated from discrete trait analysis in BEAST. B) Tree generated from structured coalescent approximation in MASCOT using BEAST2.

The results from BEAST DTA indicated the number of host type transitions was comparable between all SATs when correcting for number of taxa (Table 4). However, the SATs differed in their inferred amount of spread from wildlife to livestock and vice versa (Figure 9A-C). SAT2 had a slightly higher estimate and HPDI of wildlife to livestock transmission (µ = 1.01×10^−2^, HPDI = 7.04×10^−3^ - 2.46×10^−2^) than SATs 1 (µ = 9.49×10^−3^, HPDI = 2.60×10^−4^ - 2.29×10^−2^) and 3 (µ = 7.80×10^−3^, HPDI = 9.98×10^−5^ - 1.91×10^−2^), and notably higher estimates of livestock to wildlife transmission (µ = 6.39×10^−3^, HPDI = 4.17×10^−5^ - 1.65×10^−2^) (SAT1: µ = 8.97×10^−4^, HPDI = 4.88×10^−10^ - 3.11×10^−3^. SAT3: µ = 1.52×10^−3^, HPDI = 8.08×10^−9^ - 5.42×10^−3^) (Figure 9B). Although all SATs had a higher rate of transmission from wildlife to livestock than from livestock to wildlife, this difference was smaller in SAT2, with mean estimate for wildlife to livestock transmission within the HPDI of the estimates for livestock to wildlife transmission (Figure 9B). In contrast, the mean estimates for wildlife to livestock transmission for both SAT1 and SAT3 is clearly above the HPDI of the estimates for livestock to wildlife transmission, with the estimates for livestock to wildlife transmission showing small HPDIs and strong peaks at the smallest values (Figure 9A, 9C).

**Figure 9:**
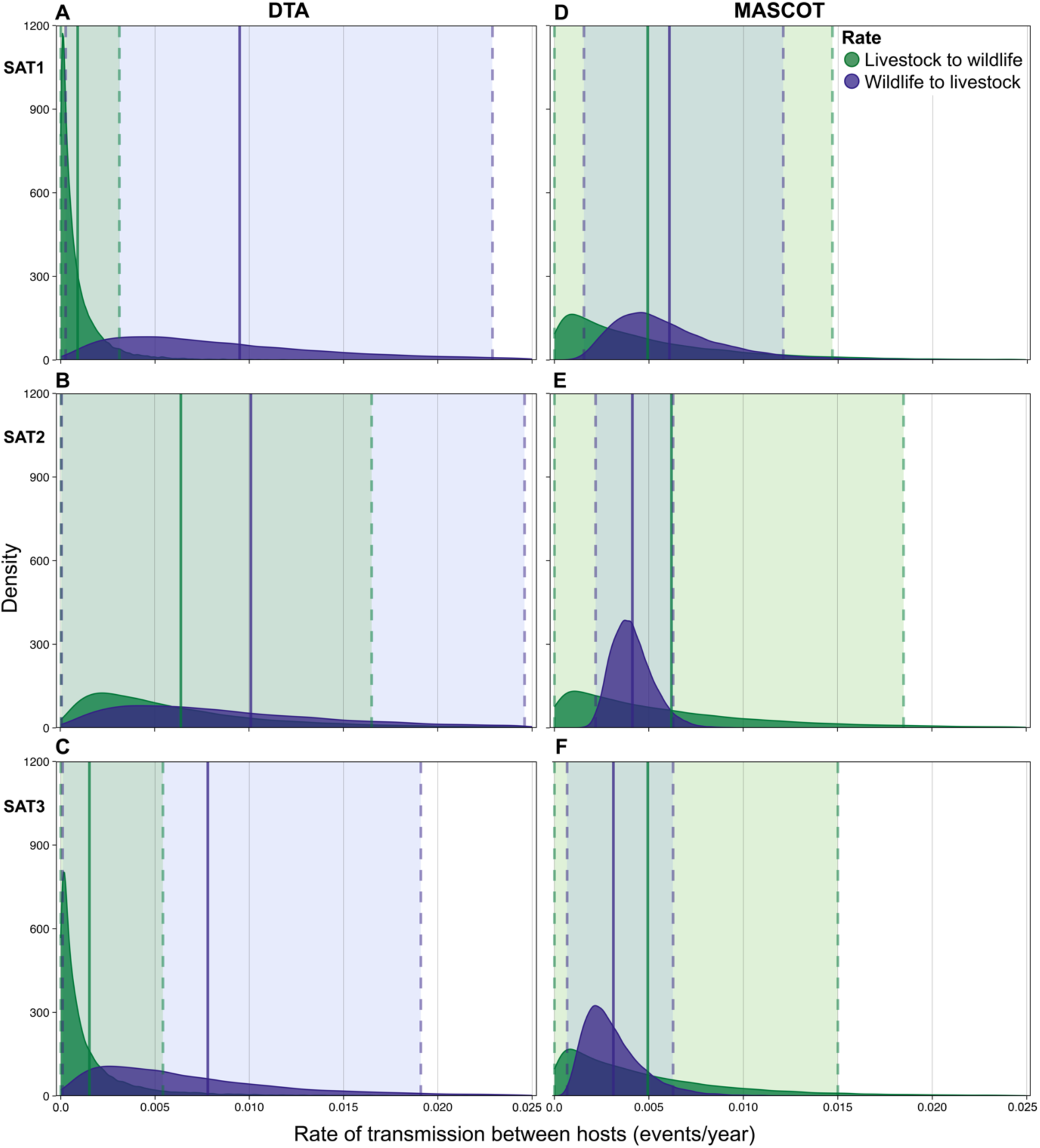
Host type transmission rates between SATs. Estimates of transmission rates from wildlife to livestock (green) and livestock to wildlife (purple) by DTA (A-C) and MASCOT (E-F) for each SAT, with all axes scaled equally. A-C)All SATs showed higher levels of wildlife to livestock transmission than livestock to wildlife transmission by DTA, with SAT2 showing higher estimates of livestock to wildlife transmission than SATs 1 and 3. E-F) All SATs showed wildlife to livestock transmission estimates farther away from 0, with estimates from MASCOT much more defined and slightly lower than estimates from DTA, and with estimates for SAT3 slightly lower than estimates for SATs 1 and 2. Estimates for livestock to wildlife transmission are more comparable for all SATs, with much larger HPDIs than wildlife to livestock transmission, or livestock to wildlife transmission calculated with DTA.

**Table 4.** **Number of estimated host type transmission events** inferred by DTA and MASCOT. Markov jump count estimations of n for each SAT are with and without unknown hosts included. Unknown hosts were included and assigned a host in DTA and excluded in MASCOT. For total migration event numbers, numbers in brackets are the estimate divided by the number of taxa (to correct for the fact that the number of inferred transmission events is expected to rise for larger numbers of samples). Met. = method, DTA = Discrete trait analysis, MAS. = MASCOT, W = Wildlife, L = Livestock.

| Parameter | Met. | SAT1 (n = 269, 252) |  | SAT2 (n = 315, 303) |  | SAT3 (n = 132, 127) |  |
| --- | --- | --- | --- | --- | --- | --- | --- |
|  |  | Estimate | HPDI | Estimate | HPDI | Estimate | HPDI |
| Migration events (all) | DTA | 35.5996 (0.132) | 33-40 | 40.957 (0.130) | 33-49 | 14.355 (0.109) | 12-19 |
|  | MAS. | 37.462 (0.148) | 36-39 | 42.156 (0.139) | 38-47 | 14.503 (0.114) | 13-17 |
| Migration events W->L | MAS. | 37.4239 | 36-39 | 41.5338 | 37-46 | 14.306 | 13-16 |
| Migration events L->W | MAS. | 0.0378 | 0-0 | 0.6223 | 0-3 | 0.1938 | 0-1 |

In contrast, inferred estimates of wildlife to livestock transmission from MASCOT were slightly lower and more precise than those from DTA. However, distributions from all SATs were still overlapping, with SAT3 slightly lower (µ = 3.12×10^−3^, HPDI = 6.69×10^−4^ - 6.28×10^−3^ than the other two (SAT1: µ = 6.08×10^−3^, HPDI = 1.56×10^−3^ - 1.21×10^−2^. SAT2: µ = 4.13×10^−3^, HPDI = 2.18×10^−3^ - 6.29×10^−3^) (Figure 9E-F). However, the estimates for livestock to wildlife transmission were different from those made with DTA, which had supported transmission in that direction, with all estimates comparable to each other and with HDPIs that extend beyond the upper estimates for wildlife to livestock transmission for all SATs (SAT1: µ = 4.93×10^−3^, HPDI = 1.18×10^−6^ - 1.47×10^−2^. SAT2: µ = 6.20×10^−3^, HPDI = 7.23×10^−7^ - 1.85×10^−2^. SAT3: µ = 4.94×10^−3^, HPDI = 1.45×10^−8^ - 1.50×10^−2^). Additionally, although the peak estimates for livestock to wildlife transmission in all SATs are lower than the peaks for wildlife to livestock transmission, these peaks are farther away from 0, and the mean estimates for livestock to wildlife transmission are above those for wildlife to livestock transmission for SATs 2 and 3 (Figure 9E-F). However, the HPDI for discrete transmission events from livestock to wildlife included 0 for all SATs, leaving the inference about this type of transmission inconclusive (Table 4).

## DISCUSSION

This study aimed to investigate whether three closely related FMDV serotypes exhibit measurable differences in molecular evolution, geographic transmission, and host taxa transmission, consistent with previously observed differences in life history. Overall, the estimates of evolutionary rate and coefficient of variation, transmission between regions, and transmission between host types suggest that the three SATs are expressing different infection patterns that are consistent with some of their observed within-host dynamics. Specifically, SAT1 was predicted to have a lower evolutionary rate due to a longer chronic carrier period (Jolles et al., 2021), and SAT2 may be transmitting more consistently from acutely infected hosts with less carrier transmission and potentially alternative maintenance hosts. These differences suggest that FMDV is evolving different strategies to persist when susceptible hosts (i.e., those who do not have immunity from past infection) are distributed inconsistently, and herds of susceptible individuals are quickly infected, recover, and become immune.

SAT1 tended to evolve slightly slower than SAT2 and SAT3, which would be consistent with SAT1 viruses spending more time in a chronic stage as predicted by previous studies (Jolles et al., 2021). This suggests that SAT1 persists in buffalo by entering a chronic carrier phase. However, the estimates for all three SATs were fairly similar to each other, varying by less than 0.001 substitutions per site per year, well within the ranges of other ssRNA virus evolution, which generally ranges from 1×10^−2^ to 1×10^−5^ substitutions per site per year, with most around 1×10^−3^ (Holmes, 2009). Our estimates of between 2.90×10^−3^ and 5.56×10^−3^ substitutions per site per year were within the same order of magnitude as several other estimates of FMDV evolutionary rates, which ranged from 1.46×10^−3^ in a whole genome analysis of all serotypes (Yoon et al., 2011), 2.369×10^−3^ in the complete open reading frame of serotype A (Mohapatra et al., 2025), and up to 6.52×10^−3^ in some non-structural regions of serotype A (Brito et al., 2018). In comparison, human T-cell lymphotropic virus II (HTLV-II) shows two orders of magnitude faster evolution depending on transmission mode (Salemi et al., 1999; Vandamme et al., 2000), which is a much larger difference than estimated for different strains of FMDV. However, HTLV-II causes persistent infection for the lifetime of the host, whereas most buffalo are thought to eventually clear their FMDV infection (Maree et al., 2016). Therefore, the effect of a carrier phase on evolutionary rates is expected to be much smaller for FMDV, and detecting any indication of a decrease in evolutionary rate in the expected direction is worth noting.

SAT3 and SAT2 were estimated to evolve at similar rates, but the estimations for SAT3 failed to converge after 200 million states, likely due to a poor clock-like signal; even though the BETS indicated that this serotype is measurably evolving, the dataset may be too small to quantify this evolution definitively. This means that the evolutionary rate presented for SAT3 may not be reliable. This study traded long sequences for a greater sample size, which provided more complete movement and host type transmission information but sacrificed the ability to reliably estimate the evolutionary rate for SAT3. A longer sequence length would help resolve SAT3’s rate of evolution and provide a stronger comparison against SAT1 and SAT2. Obtaining longer sequences might be possible if historical samples are still available for sequencing.

SAT2 exhibited much greater variation in evolutionary rate between branches compared to SAT1 and 3. We hypothesised that this pattern could be associated with host type but did not find evidence to support this; the distributions of rates for livestock and wildlife broadly overlapped. However, the distribution for wildlife contained a “wide shoulder” with more extreme values not seen in livestock, suggesting a capacity for particularly fast evolution in wildlife. Although this wide shoulder was not observed in livestock, low variation in evolutionary rate among livestock may be an artefact of sampling only during outbreaks and under disease control measures such as culling of infected livestock, which would detect only very short branches that are consistent with a small range of evolutionary rates. Therefore, it is not possible to conclude from the available data that SAT2 is consistently evolving at different rates in different host types.

Although specifics on the direction of transmission between regions varied between SATs, all SATs indicated greater movement out of Kruger National Park than in, which is consistent with Kruger being a source for FMDV to other regions, mostly to South Africa and Zimbabwe. However, this pattern is expected in this dataset, even though it includes many nodes and samples from a period of strong disease control around KNP that prevented disease transmission, especially into South Africa (Van Schalkwyk et al., 2016): When this fence was damaged during heavy flooding in 2000 and 2001, FMDV was able to re-enter previously FMDV-free South Africa, leading to KNP appearing as a source of transmission. As this dataset does not contain sequences past 2015, it cannot be assumed to represent modern transmission patterns, which may be different now that FMDV is circulating among livestock in South Africa. However, spillback from other regions into KNP and between regions not mediated by KNP suggest that transmission for all SATs in Southern Africa may have been more complex than a simple source-sink dynamic even before endemic transmission in South Africa developed.

Although there were some differences in the estimates of instantaneous rate of transmission between regions, these may be less informative than the inferred number of discrete transmission events (Markov jump counts) since the instantaneous rates are assumed to remain consistent over time. This means that rates detected now are assumed to be representative of past rates, which may not be the case. Therefore, counts of movement events may be a better quantification, and these indicate similar levels of inter-regional transmission between SATs. However, the SATs may show differences at finer spatial scales; previous work from East Africa showed waves of alternating serotypes causing outbreaks on a within-region spatial scale, suggesting that FMDV may be transmitting along several semi-overlapping disease fronts (Casey-Bryars et al., 2018). Such a pattern would not be likely to be detectable on the spatial scale used in this study, so analysing this dataset with finer-grained within-region metadata might uncover more granular differences in spatial transmission.

Interestingly, the SATs also appear to have different transmission dynamics at larger spatial scales based on the literature. SAT2 has shown the capacity to transmit long distances, causing outbreaks in North Africa and the Middle- and Near East, while SAT1 and SAT3 appear to remain mostly confined to Sub-Saharan Africa (Hall et al., 2013; WOAH and FAO, 2023). This is likely due to SAT2’s greater capacity to infect and transmit within livestock, which are far more evenly distributed than wildlife, which may be confined to refuge areas (Fana et al., 2021; Van Schalkwyk et al., 2016). Additionally, human movement of infected animals likely facilitates the long-range transmission of SAT2 (Di Nardo et al., 2025). As this serotype is the most common detected in livestock outbreaks (Fana et al., 2021), illegal movements of infected domestic animals may play a bigger role in its spatial spread than for other SATs.

The post-hoc analysis to investigate whether inter-regional transmission was driven more by livestock or by wildlife indicated no significant association between regional transmission events and host-type transmission except for in SAT1, where transmission events between wildlife were more strongly associated with inter-region transmission than expected. This analysis suggests that SAT1 cross-regional transmission events are more commonly associated with wildlife than with livestock, which is consistent with SAT1’s strong association with and maintenance within buffalo.

All SATs showed considerably more transmission from wildlife to livestock than from livestock to wildlife and this was the same across the two inference methods used (DTA and structured coalescent). These results are consistent with wildlife being the reservoir for these FMDV serotypes in southern Africa during the study period (1934-2015), which could indicate that control measures in livestock have been effective at limiting transmission. A higher rate of transmission from Kruger National Park to other regions also points to a wildlife reservoir, since the park is not inhabited by livestock and thus any spatial or host type transmission from it will be from wildlife. However, spillback from livestock into wildlife is harder to detect, as wildlife are only opportunistically sampled whereas livestock are sampled when an outbreak occurs. This, combined with the change in state from FMDV-free to FMDV-endemic in South Africa, challenges the detection of spillback from wildlife to livestock. However, this spillback may still occur. Although MASCOT did favour a set of inferred trees for SAT1 in which the roots are assigned to livestock and wildlife represent only spillover hosts, this arrangement required a much larger infected livestock population size compared to wildlife. This state is contrary to our understanding of the system; wildlife show high seroprevalence for FMDV (Thomson et al., 1992), whereas livestock infections have been relatively rare during the period covered in this study (although rates of infection in livestock have risen significantly in recent years)(Thomson, 1995; Rweyemamu et al., 2008), suggesting that the infected population of wildlife exceeds the infected population of livestock. Therefore, this result is likely an artefact of MASCOT’s methodology and does not provide strong evidence that SAT1 could be maintained by livestock.

Interestingly, the SAT2 trees generated by DTA showed several livestock to wildlife transmission events, and although these were not inferred in the MCC tree generated from MASCOT, the estimates of livestock to wildlife transmission were broad, suggesting that at least in some cases livestock may be playing a part in persistence, even if that contribution is small. This result is consistent with previous studies of SAT2 viruses, which suggest that transmission within cattle and from cattle to wildlife do occur (Hall et al., 2013; Brito et al., 2016). Transmission from livestock to wildlife may also be part of the infection strategy for some strains of SAT2, which is a more common serotype in cattle and thought to cause fewer chronic infections (Fana et al., 2021; Jolles et al., 2021), and thus might require a larger critical community size. As transmission from livestock back to wildlife could make the metapopulation of susceptible hosts larger, thus supporting a pathogen needing larger populations to persist (Viana et al., 2014).

These results also suggest that FMDV containment efforts have been less complete for SAT2 than for SATs 1 and 3. Indeed, SAT2 causes more detected outbreaks among both cattle and wildlife than either SAT1 or SAT3, also suggesting that its dynamics are different (Bastos et al., 2003; Dyason, 2010; Fana et al., 2021; Hall et al., 2013). This is consistent with this study’s findings; both SAT2 trees showed several instances of sustained transmission among cattle over a short period of time, whereas the trees for SAT1 and SAT3 do not. This could occur if SAT2 were able to spread more easily in cattle before the infection is noticed and control measures implemented, or if it is better adapted to persistence in cattle in other ways. Alternatively, increased environmental persistence could lead to greater transmission from livestock to wildlife, particularly in the case of wildlife grazing on pastures recently inhabited by livestock and containing fresh manure, since FMDV type O has been shown to persist in bovine manure for several days (Colenutt et al., 2020). Such adaptations would be consistent with a different life history strategy.

Overall, this study suggests that the three SATs do indeed show phylodynamic patterns that are consistent with different life history strategies, including patterns of evolution that are consistent with chronic infections and transmission between host types being more frequent in some serotypes than others. However, these findings pose further questions. For example, the representative strain of SAT1 studied by Jolles et al. (2021) had a higher R_0_ estimate in buffalo compared to the other SATs, even without including carrier transmission, which is inconsistent with the greater number of outbreaks and long-range transmission seen in SAT2 (Bastos et al., 2003; Dyason, 2010; Hall et al., 2013). The fact that we see evidence of more dynamic infection among SAT2, including potential transmission from livestock to wildlife, suggests that other life history strategy differences are affecting FMDV epidemiology. Particularly, the three serotypes likely have different dynamics in cattle and in buffalo: Buffalo usually show substantially fewer clinical signs of FMD than cattle do, and differences in pathogenesis may lead to differences in transmission both across FMDV strains and among host species (Maree et al., 2016; Stenfeldt et al., 2025). Additionally, increased levels of environmental persistence may contribute to some of these dynamics (Colenutt et al., 2020), such as the potential transmission of SAT2 from livestock back to wildlife. Different segments of the genome may also provide novel information. This study focused exclusively on VP1 sequences, but FMDV evolutionary rate estimates can vary within the genome (Brito et al., 2018), and recombination in these regions is also possible (Cortey et al., 2019). Analyses of whole genomes would provide more detailed and textured reconstructions of FMDV evolutionary and transmission dynamics.

Different successful life history strategies among FMDV serotypes, and likely also among strains within these serotypes, illustrate that there are many ways to be an effective virus. These patterns we observed may suggest between-pathogen competition selecting for different viral phenotypes and strategies. Within-host competition between SATs has already been observed: SAT1 out-competes SAT2 and SAT3 when co-infected in cell culture, causing elimination of SAT2 and SAT3 (Maree et al., 2016). Virus competition is known to affect dynamics in other viruses as well with mechanisms other than cross-immunity (Isaacs and Burke, 1959; Folimonova, 2012; Dee et al., 2021), causing significant effects in virus occurrence (Nickbakhsh et al., 2019). The presence of competitive interactions between SATs shows that the buffalo-FMDV system has plenty of scope for further fundamental research on genotype-phenotype relationships and on the traits that allow viruses to persist and co-exist. Further work should aim to reveal how these traits connect to their molecular evolution, and the mechanisms that might drive these adaptions, offering insights that have both theoretical and practical applications to disease management, virology, ecology, and epidemiology.

## Acknowledgments

The authors thank Melanie Chitray for comments on an earlier draft.

## Author contributions

**Avery Holmes:** Conceptualization, Data curation, Formal analysis, Investigation, Methodology, Visualization, Writing - original draft, Writing - review & editing.

**Eva Pérez-Martin:** Conceptualization, Writing - review & editing

**Simon Gubbins:** Conceptualization, Funding acquisition, Resources, Writing - review & editing.

**Brianna Beechler:** Conceptualization, Funding acquisition, Writing - review & editing.

**Anna Jolles:** Conceptualization, Funding acquisition, Writing - review & editing.

**Roman Biek:** Conceptualization, Funding acquisition, Methodology, Project administration, Resources, Supervision, Writing - review & editing.

