## Supplementary materials for "Evolutionary analysis supports variation in life history strategies between three foot-and-mouth-disease-virus serotypes"

***Table S1: TempEst results** for each SAT.*

| Clade | Number of Sequences | Date range | Evolutionary rate | $R^2$ |
|---|---|---|---|---|
| SAT1 Full tree | 269 | 1961-2015 (54 years) | 6.1681E-4 | 1.3924E-2 |
| SAT1 Clade 1 | 152 | 1966-2015 (49 years) | 1.206E-3 | 0.1172 |
| SAT1 Clade 2 | 72 | 1977-2003 (26 years) | 2.5993E-3 | 0.1707 |
| SAT1 Clade 3 | 43 | 1961-2006 (45 years) | 1.1507E-3 | 0.2468 |
| SAT2 Full tree | 315 | 1948-2015 (67 years) | 1.7493E-3 | 0.1458 |
| SAT2 Clade 1 | 197 | 1948-2015 (67 years) | 1.4467E-3 | 0.1112 |
| SAT2 Clade 2 | 96 | 1969-2015 (46 years) | 4.5349E-4 | 3.1645E-4 |
| SAT2 Clade 1+2 | 293 | 1948-2015 (67 years) | 1.2541E-2 | 9.9021E-2 |
| SAT3 Full tree | 132 | 1934-2010 (76 years) | 1.0336E-4 | 4.5926E-4 |
| SAT3 Clade 1 | 74 | 1969-2010 (41 years) | 2.5078E-4 | 7.6459E-3 |
| SAT3 Clade 2 | 29 | 1990-1991 (1 year) | -0.0181 | 0.2062 |
| SAT3 Clade 3 | 23 | 1961-1998 (37 years) | -0.0005 | 7.9973E-2 |

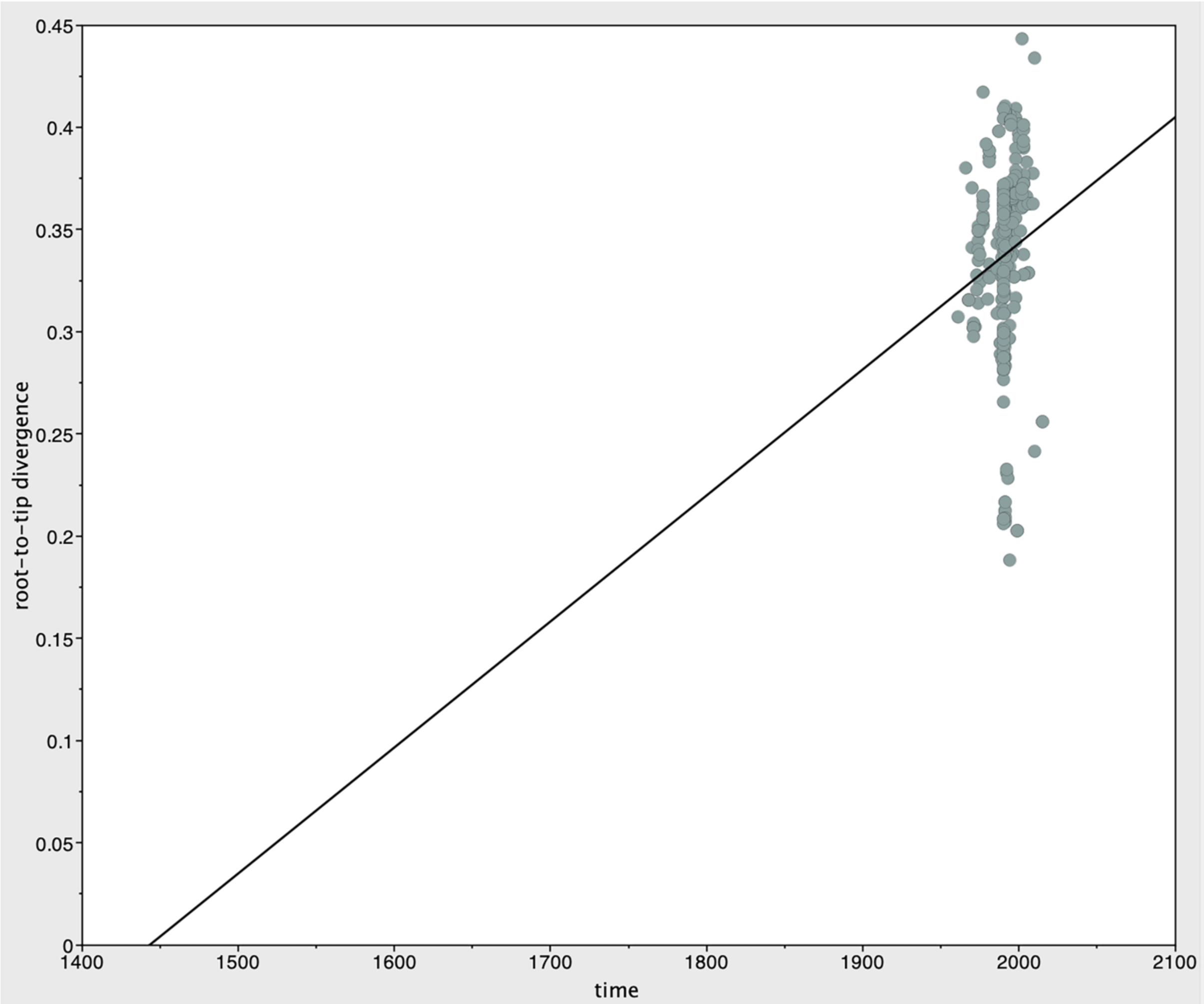

***Figure S1: TempEst regression graph for SAT1, full tree,*** *consistent with a molecular clock signal.*

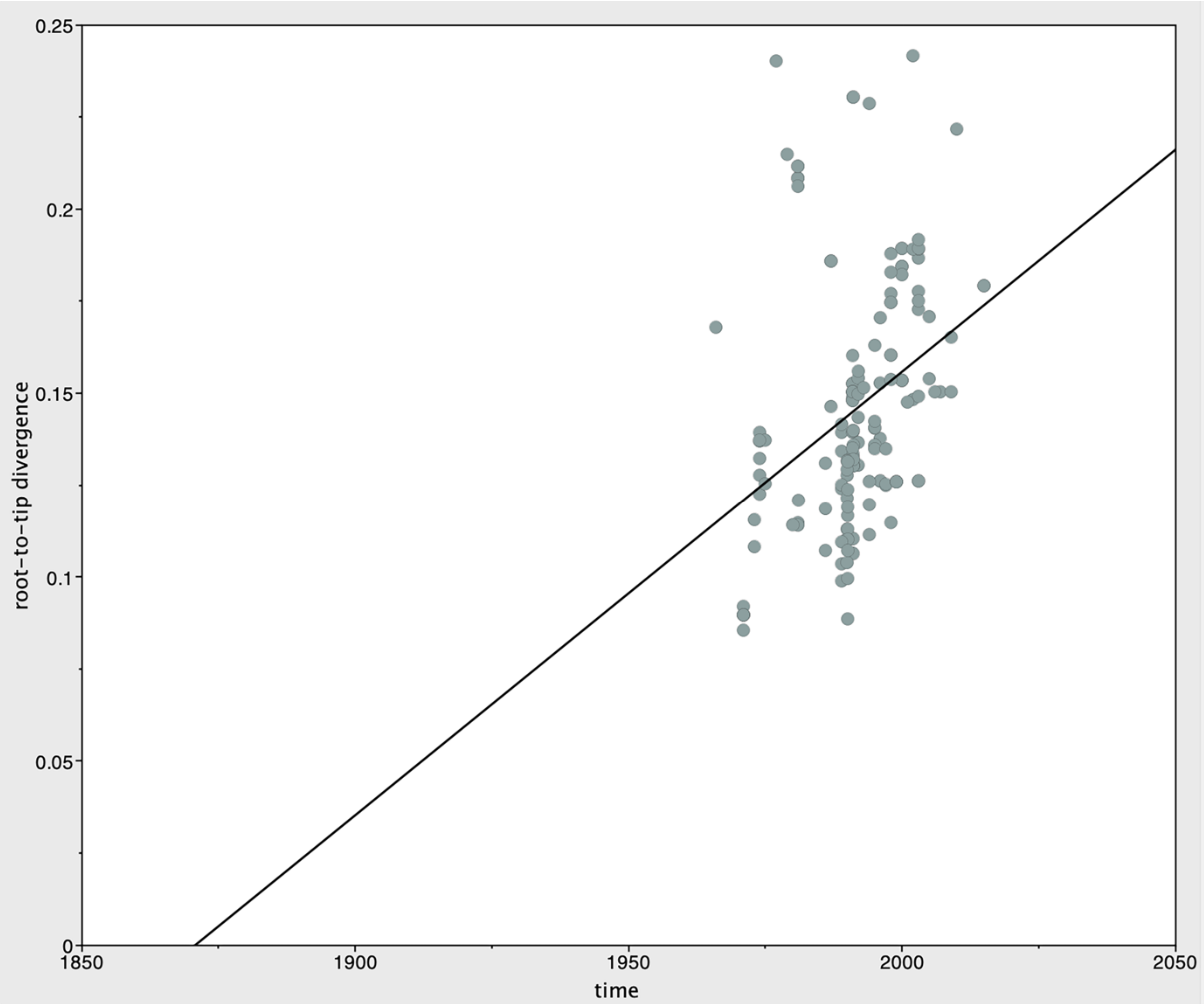


***Figure S2: TempEst regression graph for SAT1, subtree 1,*** *consistent with a molecular clock signal.*

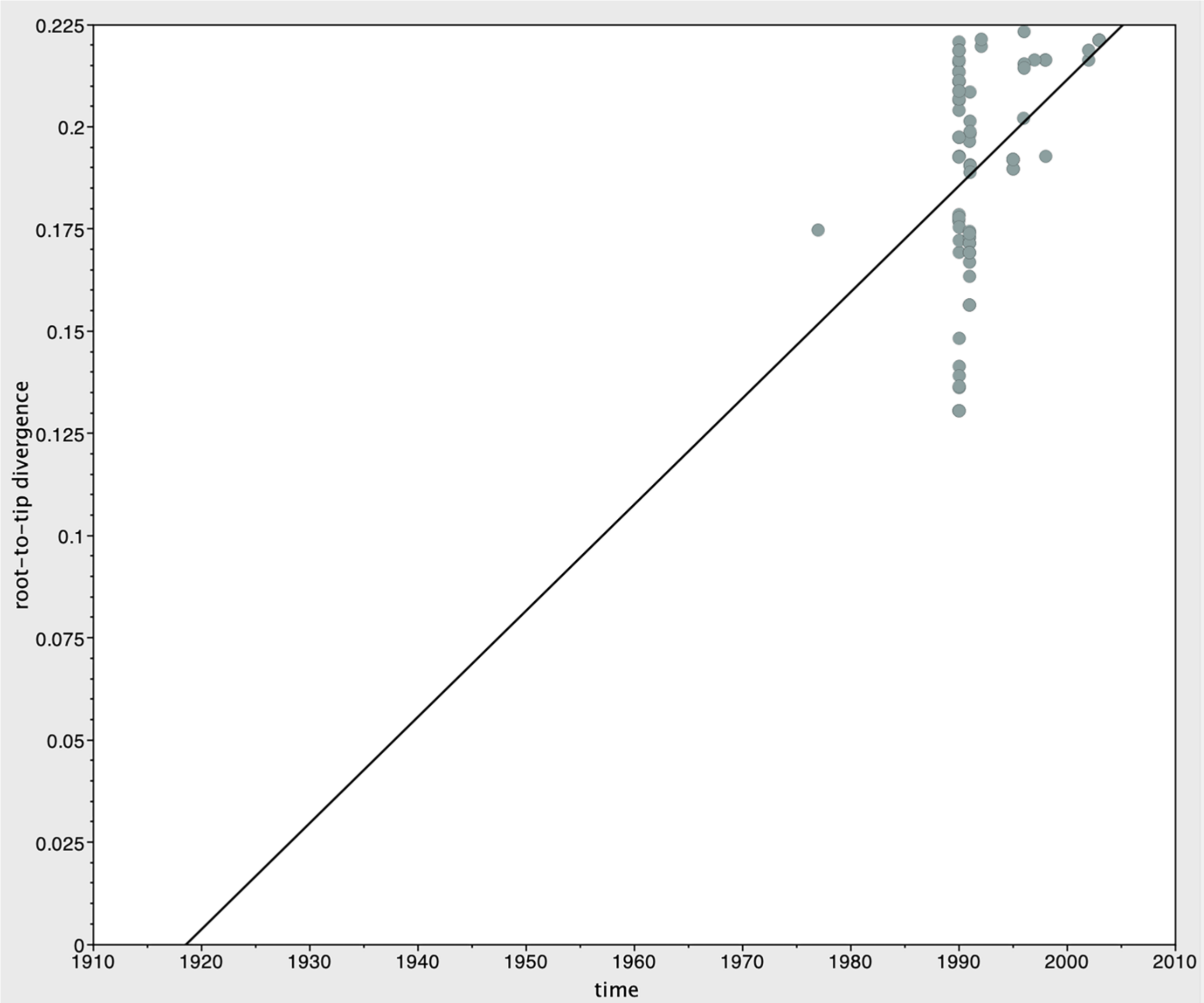


***Figure S3: TempEst regression graph for SAT1, subtree 2,*** *indicating a clear molecular clock signal.*

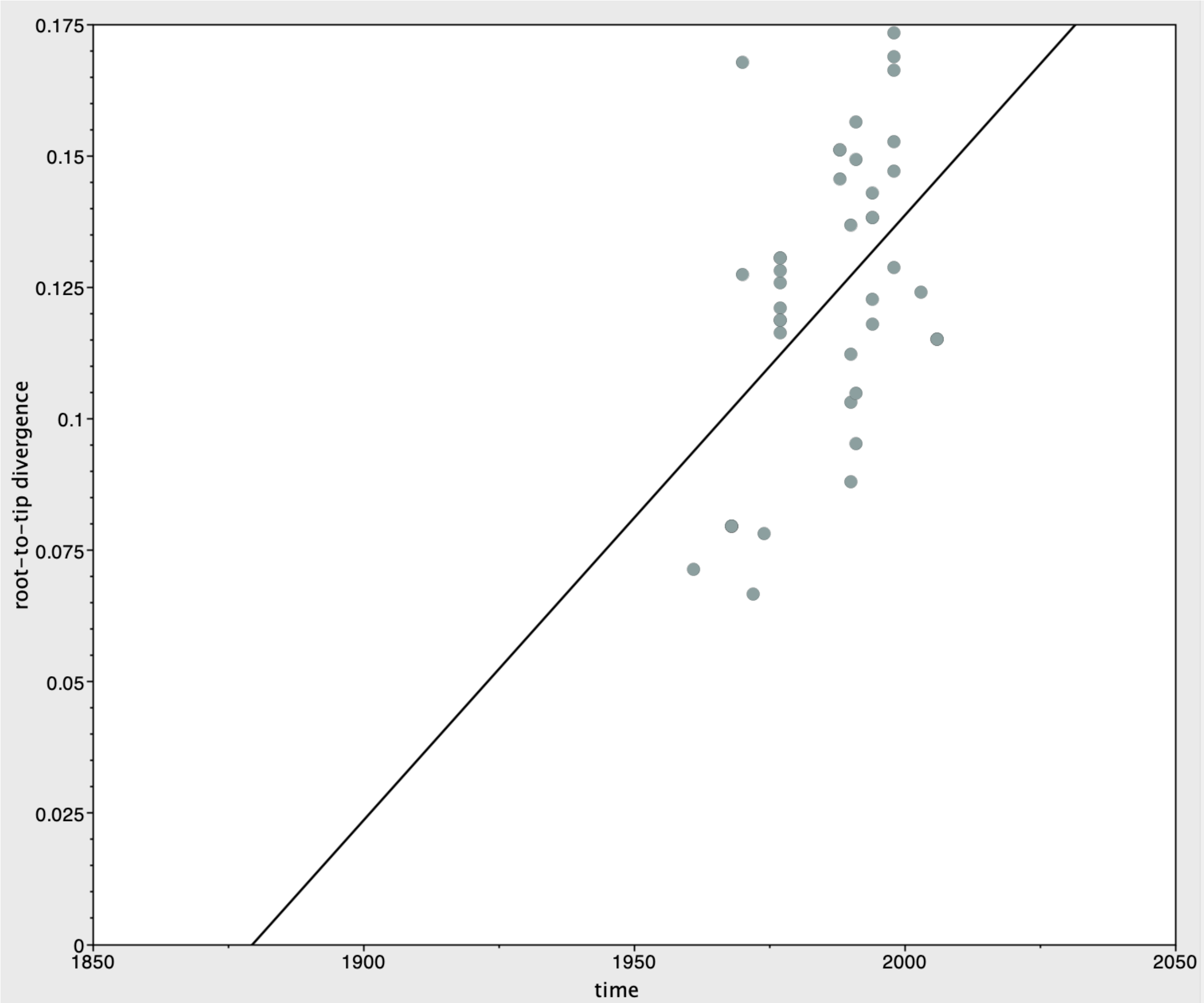


***Figure S4: TempEst regression graph for SAT1, subtree 3,*** *consistent with a molecular clock signal.*

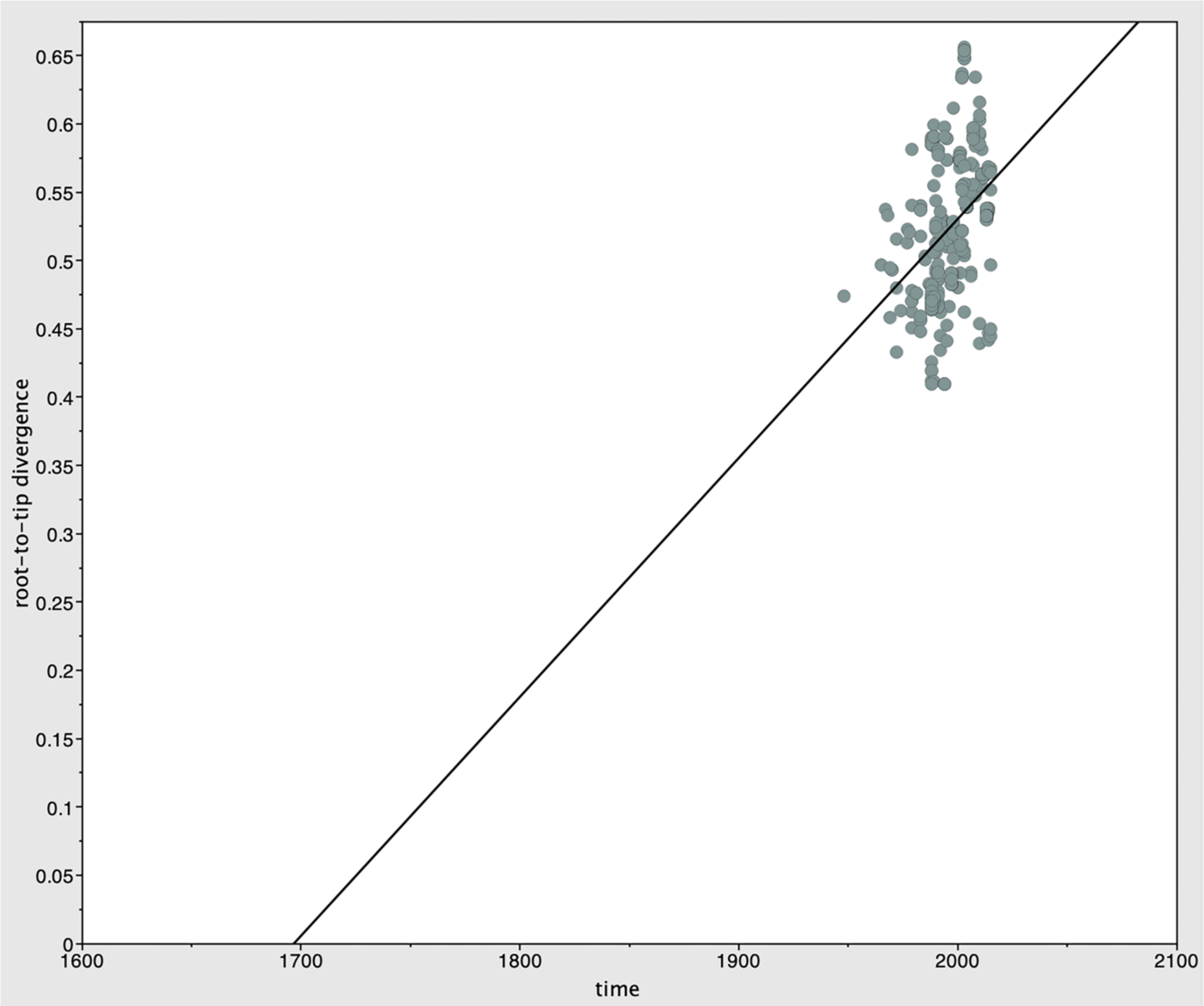


***Figure S5: TempEst regression graph for SAT2, full tree,*** *consistent with a molecular clock signal.*

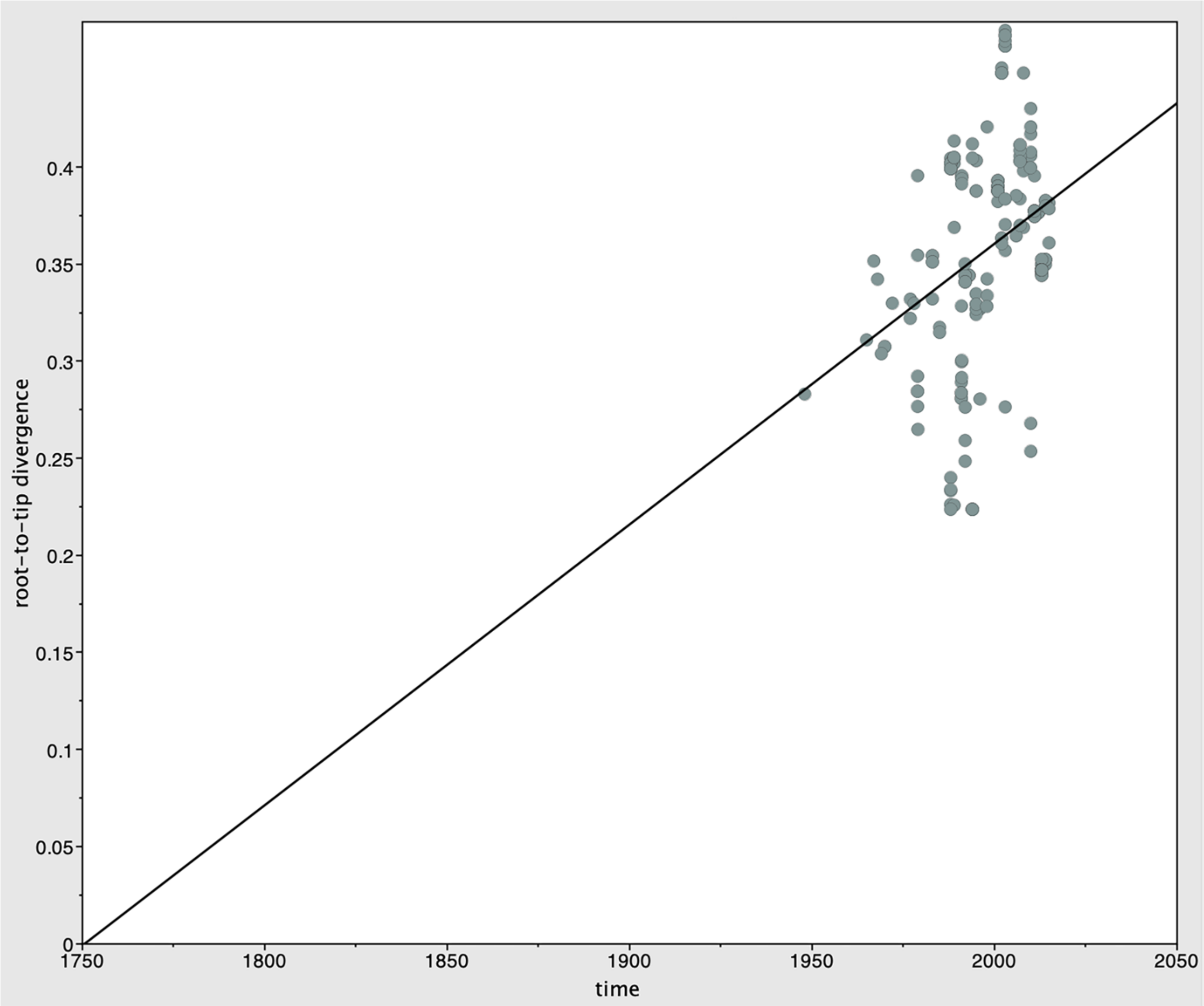


***Figure S6: TempEst regression graph for SAT2, subtree 1,*** *consistent with a molecular clock signal.*

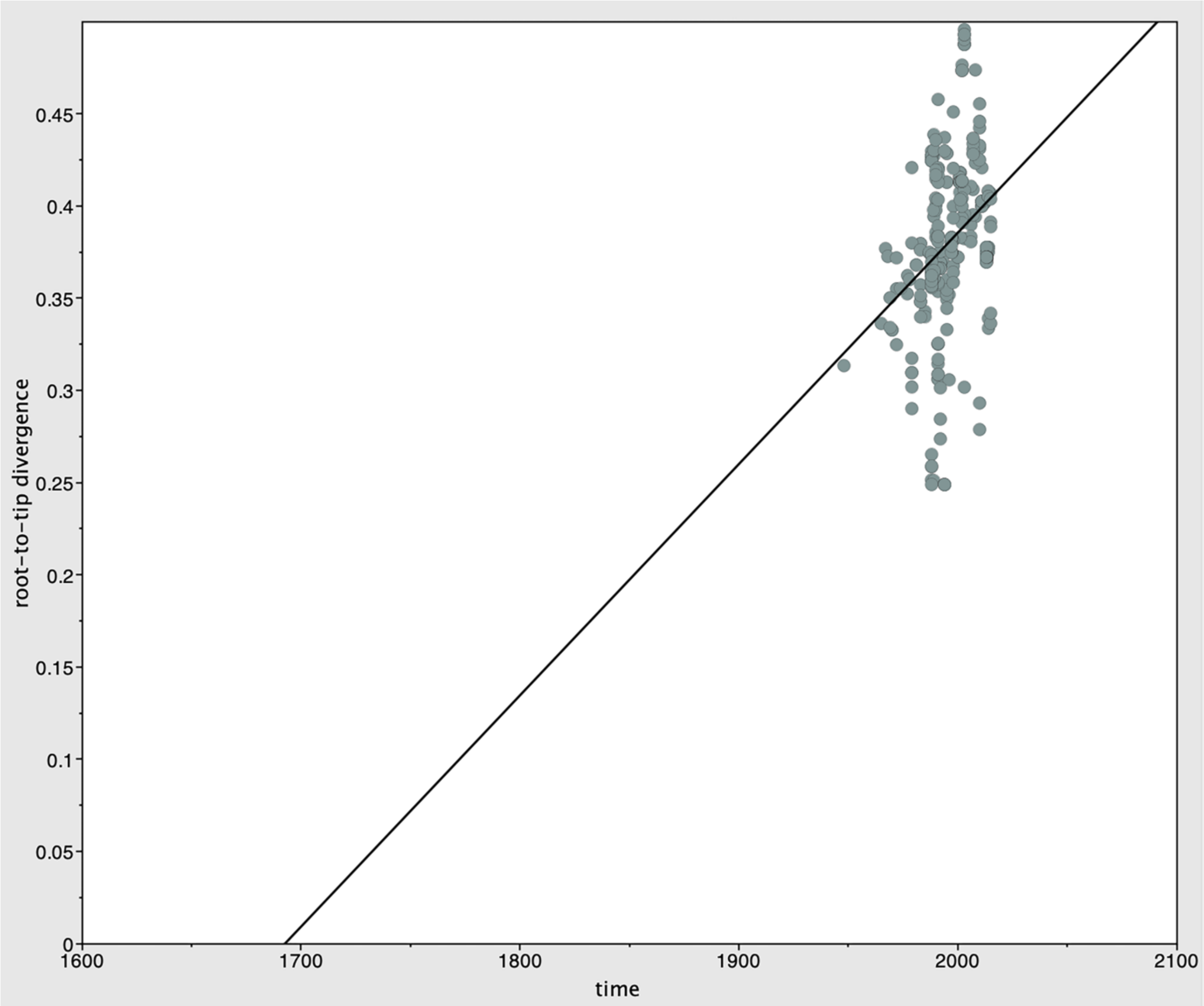


***Figure S7: TempEst regression graph for SAT2, subtrees 1 and 2 combined,*** *consistent with a molecular clock signal.*

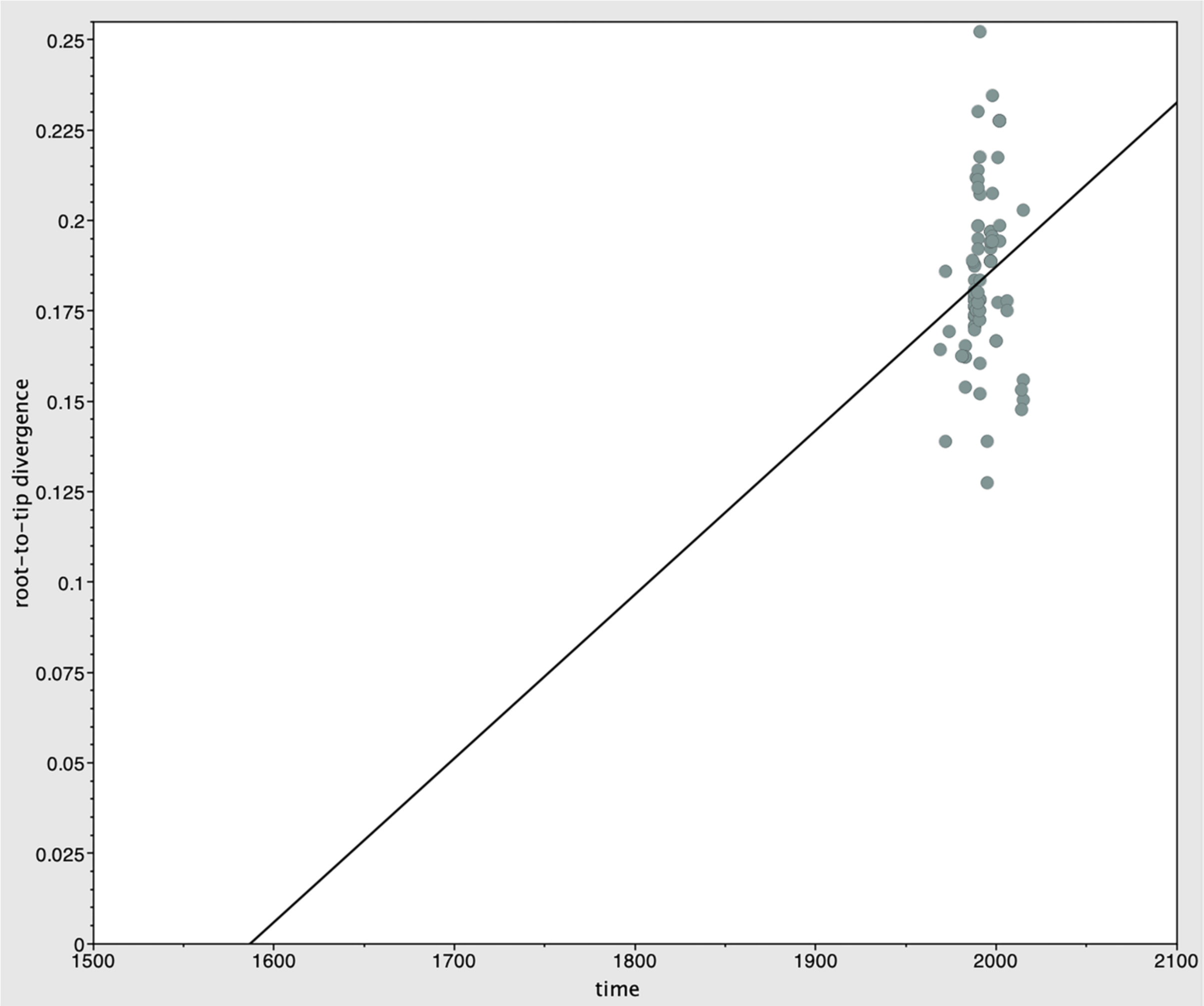


***Figure S8: TempEst regression graph for SAT2, subtree 2,*** *consistent with a molecular clock signal.*

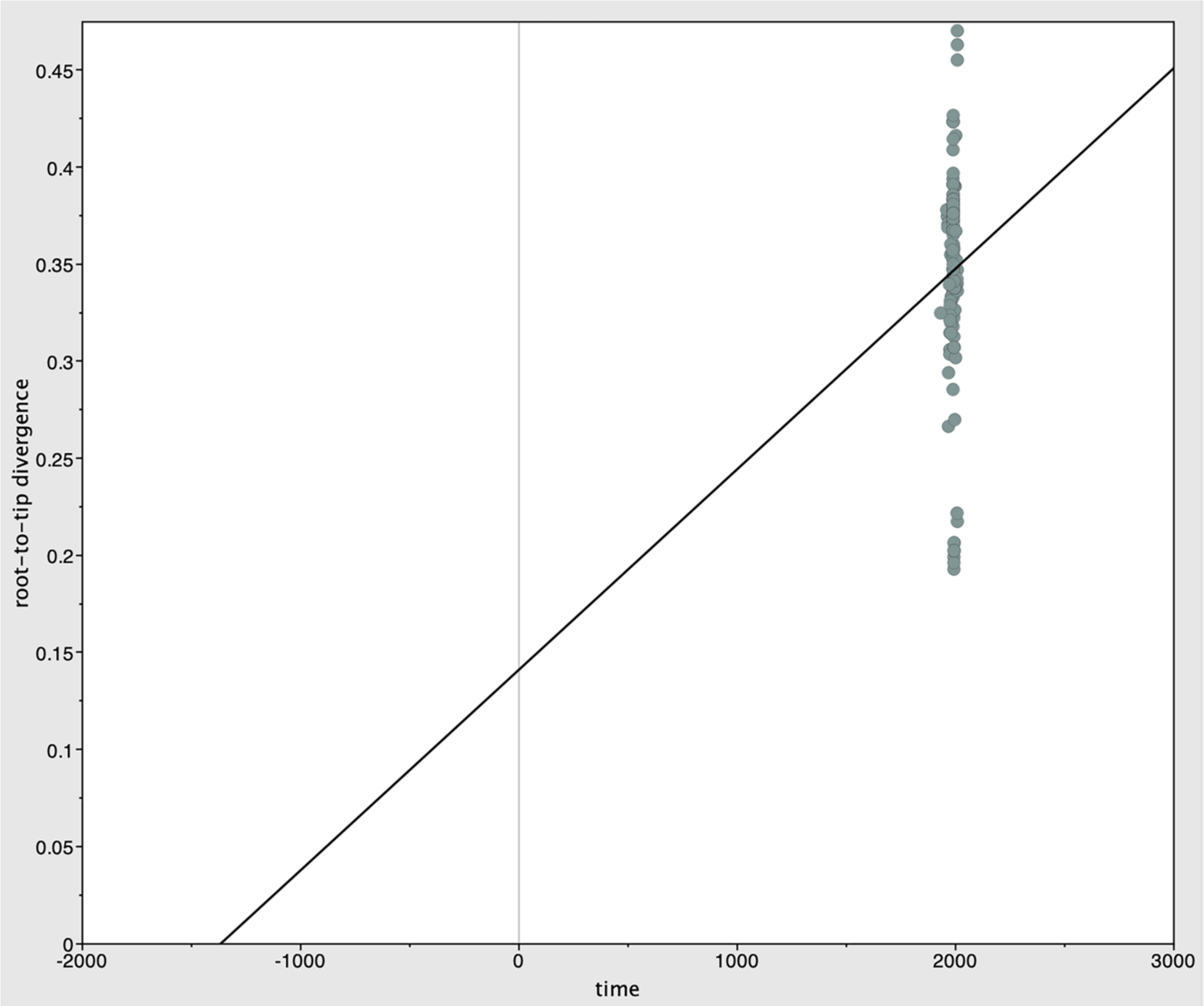


***Figure S9: TempEst regression graph for SAT3, full tree,*** *suggesting a weak molecular clock signal, although there is considerable variation in the data, suggesting that the signal may be unreliable.*

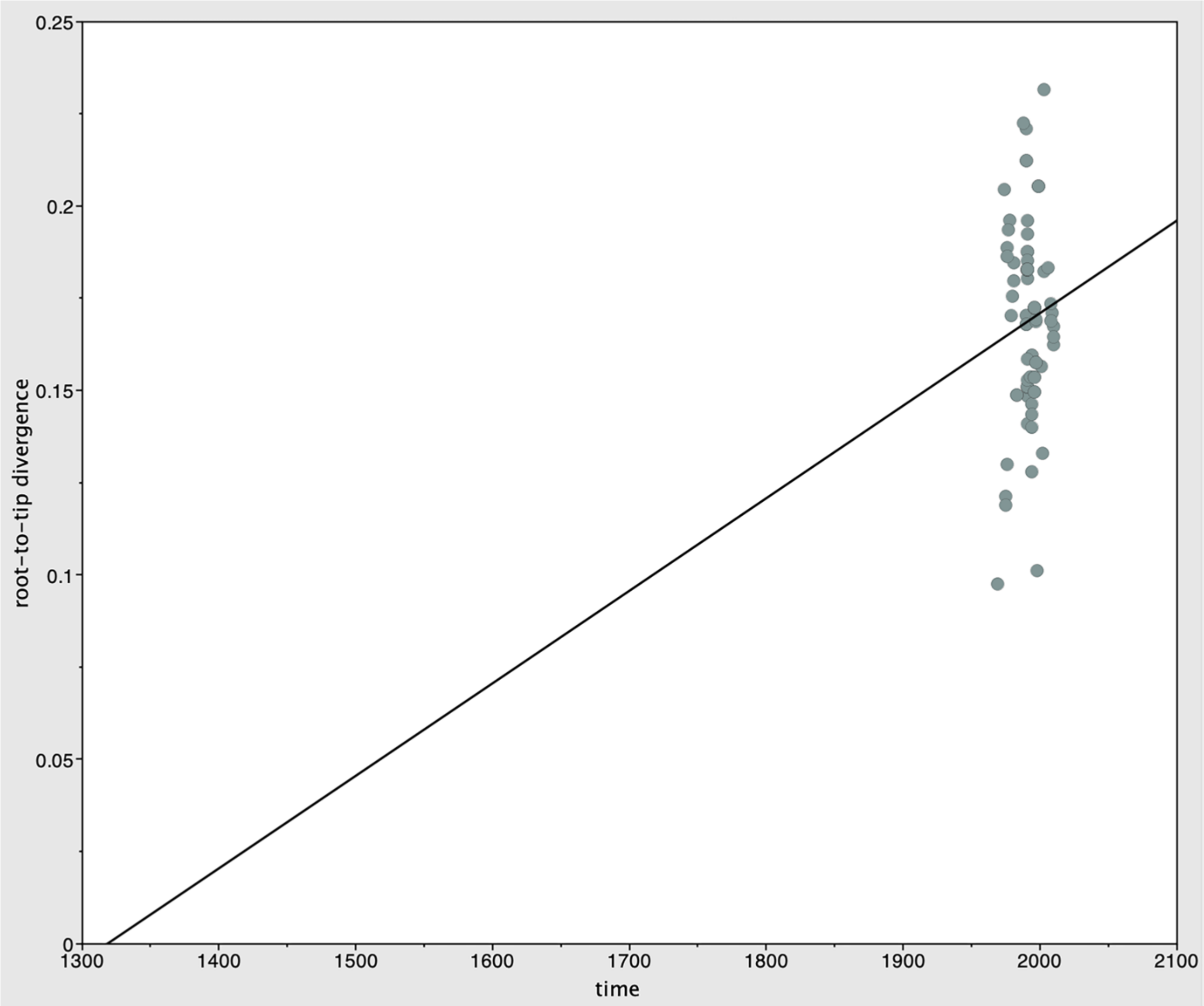


***Figure S10: TempEst regression graph for SAT3, subtree 1,*** *consistent with a molecular clock signal.*

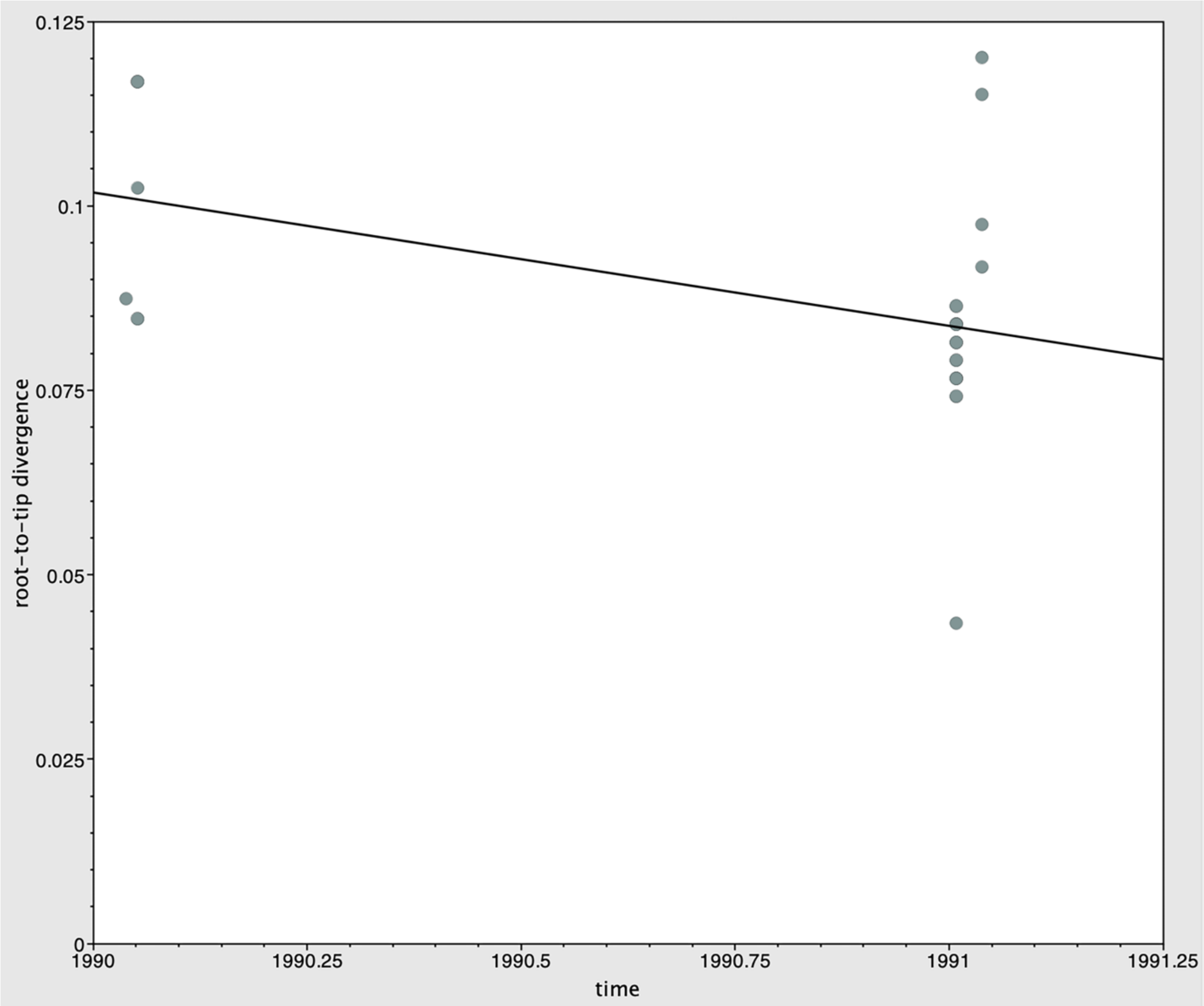


***Figure S11: TempEst regression graph for SAT3, subtree 2,*** *indicating a negative molecular clock signal, which indicates that no molecular clock signal is detectible in this clade.*

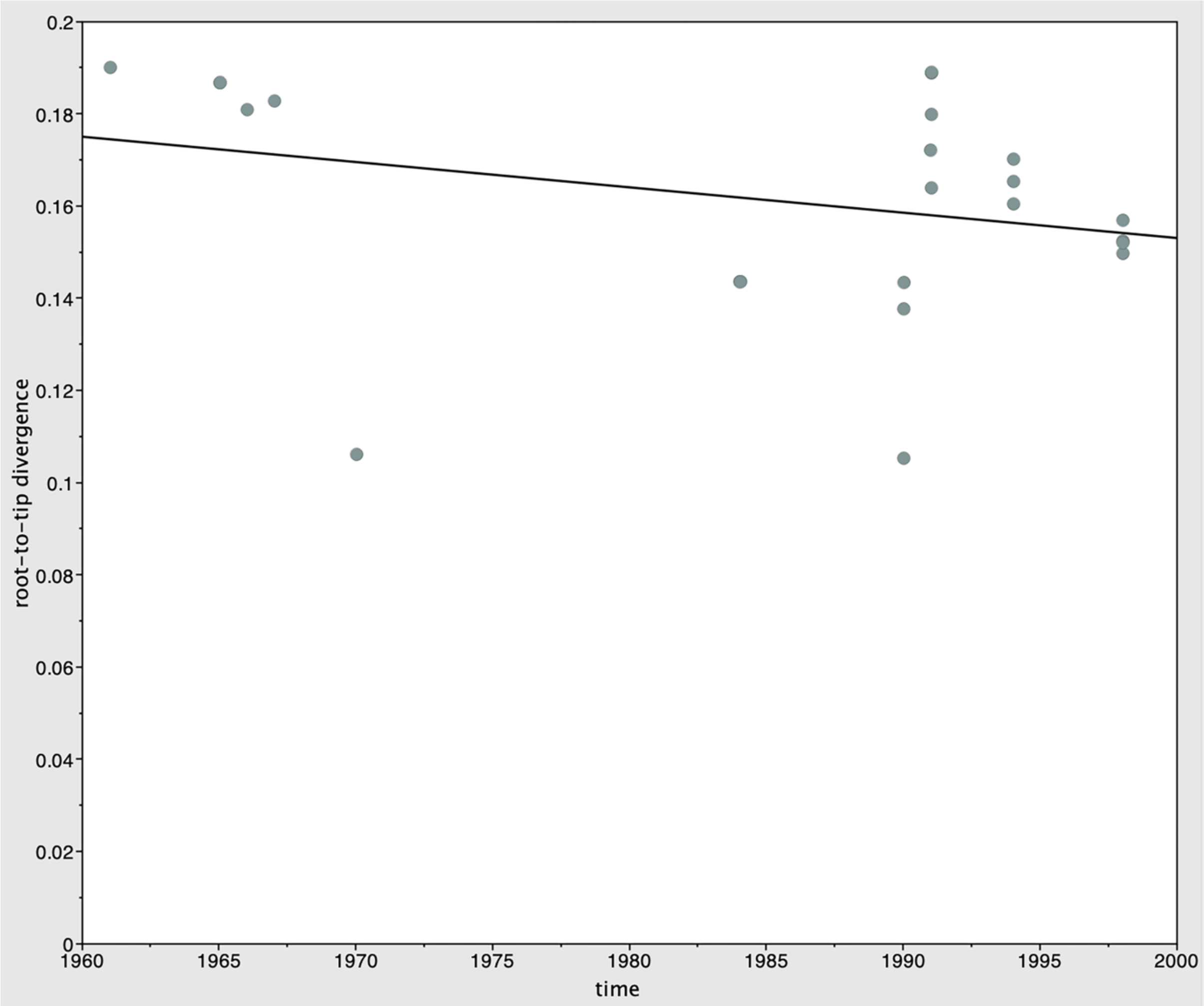


***Figure S12: TempEst regression graph for SAT3, subtree 3,*** *indicating a negative molecular clock signal, which indicates that no molecular clock signal is detectible in this clade.*

***Table S2: Bayesian evaluation of temporal signal results.*** *Models without tip dates (isochronous) and with a fixed clock rate were compared against models with tip dates included (heterochronous) and a relaxed clock. A Bayes factor test indicated that the heterochronous model was preferred for all subtrees in all SATs (values over 3 indicate strong support, values over 5 indicate very strong support (Duchene et al., 2020)).*

| SAT | Subtree | Tips | Clock rate | Log likelihood | Bayes factor |
|---|---|---|---|---|---|

| | | | | | |
|---|---|---|---|---|---|
| 1 | 1 | Heterochronous<br>Isochronous | Relaxed<br>Fixed | -9106.73<br>-9172.33 | 65.60 |
| 1 | 2 | Heterochronous<br>Isochronous | Relaxed<br>Fixed | -4257.20<br>-4261.90 | 4.70 |
| 1 | 3 | Heterochronous<br>Isochronous | Relaxed<br>Fixed | -3142.28<br>-3159.10 | 16.82 |
| 2 | 1 | Heterochronous<br>Isochronous | Relaxed<br>Fixed | -10022.27<br>-10193.06 | 170.79 |
| 2 | 1+2 | Heterochronous<br>Isochronous | Relaxed<br>Fixed | -14373.73<br>-14620.85 | 247.12 |
| 2 | 2 | Heterochronous<br>Isochronous | Relaxed<br>Fixed | -4580.90<br>-4651.25 | 70.36 |
| 3 | 1 | Heterochronous<br>Isochronous | Relaxed | -5288.84<br>-5319.41 | 30.57 |
| 3 | 2 | Heterochronous<br>Isochronous | Relaxed<br>Fixed | -1461.91<br>-1472.64 | 10.72 |
| 3 | 3 | Heterochronous<br>Isochronous | Relaxed<br>Fixed | -2468.54<br>-2473.61 | 5.07 |

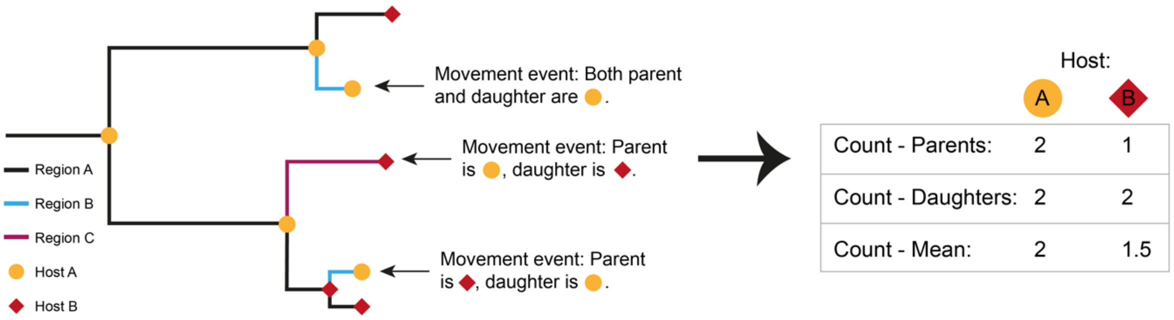


***Figure S13: Simplified diagram of regional movement host association counting method**. All regional movements (changes in colour in the branches) were identified, and the host associated with the event was assigned according to the identity of the parent or the daughter. The total number of movement events was calculated once each for both the parent and daughter node counts, and then the mean was taken between these two values.*
